# Glucose hypometabolism and hyperphosphorylated Tau synergistically drive neuronal necroptosis

**DOI:** 10.64898/2026.02.01.703076

**Authors:** Xiaoshi Chen, Sixuan Li, Antonia Neubauer, Xiang Li, Wei Mao, Matthias Brendel, Jixuan Liu, Mengmeng Zhang, Chongzhe Yang, Ruisheng Xu, Jianpin Liu, Bin Shan, Junying Yuan, Cong Liu, Jochen Herms, Chengyu Zou

## Abstract

The combination of brain glucose hypometabolism and hyperphosphorylated Tau (p-Tau) pathology is the strongest known clinical predictor of imminent cognitive decline, yet how these factors cooperate to drive dementia remains unknown. Here, we show that glucose hypometabolism synergizes with p-Tau to trigger neuronal loss through necroptosis. Under low-glucose conditions, accumulated p-Tau forms a molecular scaffold that directly recruits the necroptotic kinase RIPK1, while concomitant loss of the necroptosis checkpoint protein A20 removes a critical brake on this death pathway. This dual mechanism thereby precipitates neuronal necroptosis independent of classical TNF/TNFR1 signaling paradigm. Restoring A20 expression with acetyl-L-carnitine as a dietary supplement or preventing the p-Tau – RIPK1 interaction using a RIPK1 derived competitive peptide alleviates neuronal necroptosis and brain atrophy in a Tau transgenic mouse model. Collectively, our findings uncover a previously unrecognized metabolism-driven necroptotic signaling cascade initiated by a p-Tau-RIPK1 hub, providing mechanical insight into how glucose hypometabolism synergizes with p-Tau to drive neurodegeneration.

## INTRODUCTION

Tauopathies, including Alzheimer’s disease (AD), frontotemporal dementia (FTD) and progressive supranuclear palsy (PSP), are neurodegenerative disorders characterized by the abnormal accumulation of hyperphosphorylated Tau (p-Tau) protein in neurons. While p-Tau has been recognized as a key determinant of neuronal loss, dementia progression and severity, its pathology typically appears decades before clinical symptoms arise^1,2^. This prolonged preclinical phase suggests that p-Tau accumulation precedes neurodegeneration and raises the possibility that additional pathological cofactors may be required to trigger neuronal loss.

Recent studies have proposed a link between p-Tau and necroptosis in driving neuronal loss and cognitive decline in AD^3–5^. However, given the long latency between p-Tau pathology and cognitive symptom onset, the factor(s) that orchestrate the progression from p-Tau accumulation to necroptotic cell death in vivo is unclear. Identifying the bridging factor(s) is essential for understanding disease progression and developing effective therapeutic strategies.

In addition to p-Tau pathology, brain glucose hypometabolism, characterized by reduced cerebral glucose uptake, is a well-recognized clinical feature of neurodegenerative diseases^6,7^. Among diagnostic biomarkers for dementia, glucose hypometabolism patterns detected by FDG-PET provide the highest predictive accuracy in distinguishing individuals with mild cognitive impairment (MCI) who would soon progress to AD or FTD dementia from those who would remain stable or revert to a normal cognitive state^8,9^. Growing evidence indicates that glucose hypometabolism not only precedes neurodegeneration but also may play an active role in driving decline of neurological functions^10–12^. Moreover, interventions that enhance cerebral glucose availability have shown promise in improving memory function in p-Tau associated pathologies^13,14^. Despite these insights, the mechanistic contribution of glucose hypometabolism to neuronal loss in tauopathies remains poorly understood.

Necroptosis has been established as a TNF receptor-1 (TNFR1) signaling dependent paradigm that promotes necrotic cell death under caspase-deficient conditions^15,16^. Upon engagement by TNFα (TNF), TNFR1 organizes the sequential formation of signaling complexes to execute apoptosis and necroptosis. The interaction of kinase activated RIPK1 with RIPK3 and MLKL leads to the formation of Complex IIb which in turn executes necroptosis^15,17^. Necroptosis is tightly regulated by a sophisticated network of molecular checkpoints, including the ubiquitin-editing enzyme A20 (also known as tumor necrosis factor induced protein 3, TNFAIP3) which suppresses necroptosis by targeting both RIPK1 and RIPK3^18,19^. How RIPK1 and necroptosis are activated outside the canonical TNFR1 signaling pathway has yet to be elucidated.

This study reveals that glucose hypometabolism acts in concert with p-Tau to drive neuronal necroptosis. Our findings demonstrate that under low-glucose conditions, p-Tau accumulation forms a molecular platform that directly recruits and activates RIPK1, independent of TNF/TNFR1 signaling. The downregulation of the necroptotic checkpoint A20 under glucose hypometabolism removes a critical brake on RIPK1 kinase activity dependent necroptosis. We further show that reduced glucose uptake coincides with downregulated A20 expression and increased RIPK1 activation in hippocampal subregions affected by p-Tau pathology in AD patients.

Taken together, these results indicate that glucose hypometabolism might drive neurodegeneration through a double-hit mechanism independent of classical TNF/TNFR1 signaling paradigm, by promoting the formation of a p-Tau–RIPK1 signaling hub that initiates necroptosis and by downregulating the necroptotic checkpoint A20. Our study provides a new paradigm for the mechanism of necroptosis and shed new mechanistic light on the intracellular pathways linking glucose hypometabolism to necroptotic activation in tauopathies.

## RESULTS

### Glucose hypometabolism induces necroptosis in neurons harboring p-Tau

The human P301S Tau mutation leads to the formation of filamentous pathology composed of p-Tau^20^. To investigate the mechanisms underlying p-Tau induced neuronal loss, we generated a neuronal cell line stably expressing the P301S mutant Tau protein (P301S-Tau HT22) (Figure S1A). P301S-Tau expressing HT22 cells were cultured in high-glucose DMEM (25 mM glucose) according to a standard protocol^21^. P301S-Tau expressing HT22 cells exhibited no evidence of cell death under these culture conditions. The basal concentration of extracellular glucose in the brain is 1.5 mM, substantially lower than the concentration in the medium used for cell culture^22^. We further observed that P301S-Tau HT22 cells remained alive when cultured in medium with a physiological glucose concentration of 1.5 mM for 48 hours (Figure 1A).

**Figure 1.**
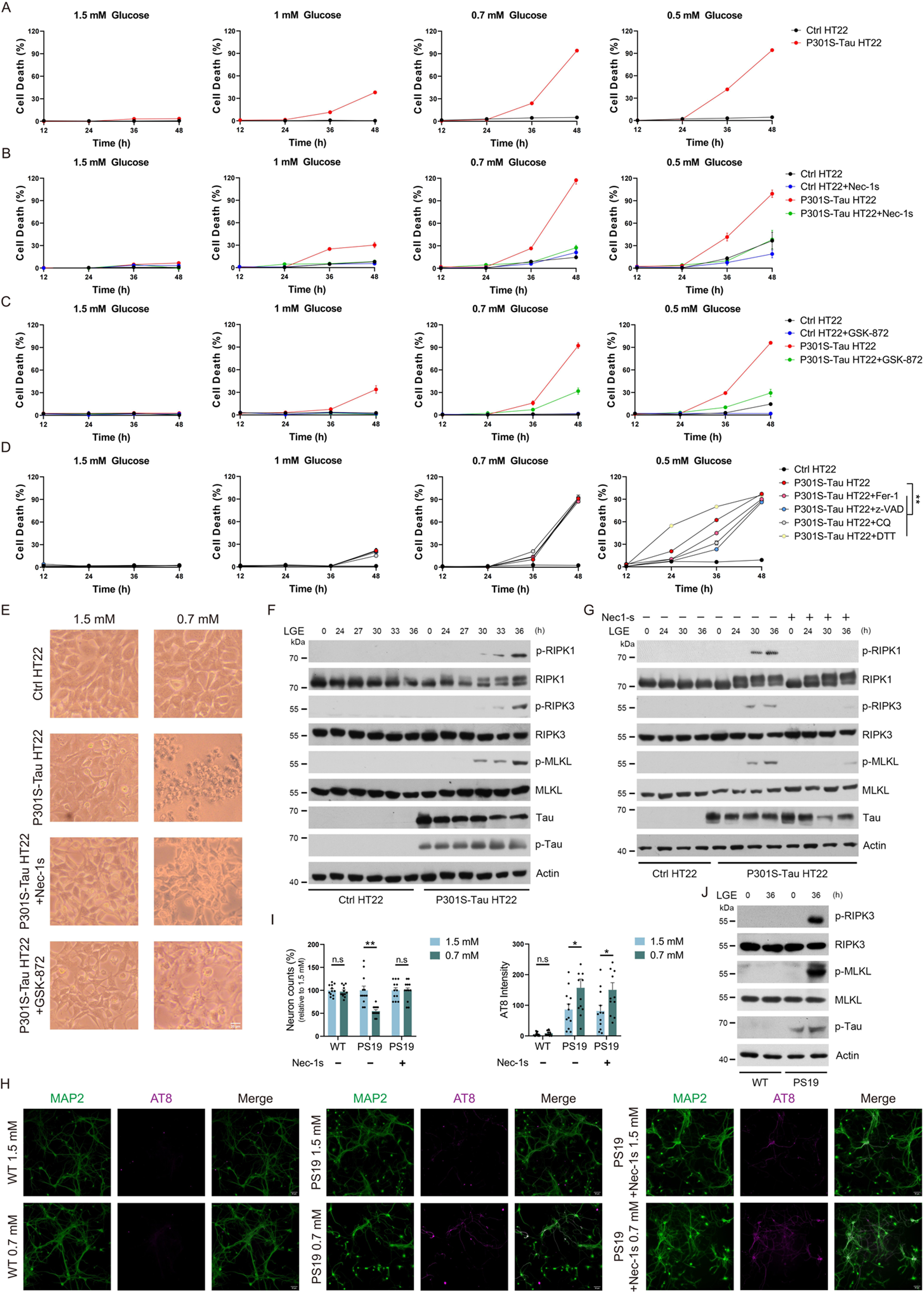
Glucose hypometabolism triggers necroptosis in neurons harboring p-Tau. (A-C) HT22 cells stably expressing the P301S mutant human Tau protein (P301S-Tau HT22) or control vector (Ctrl HT22) were cultured in medium with specified glucose concentrations for up to 48 hours (A). Where indicated, the culture medium was supplemented with either the RIPK1 inhibitor Nec-1s (10 µM, diluted 1:2000 from DMSO stock), the RIPK3 inhibitor GSK-872 (1 µM, diluted 1:5000 from DMSO stock), or DMSO alone as controls at the corresponding concentrations (B-C). Cell death was assessed using SYTOX Green (SG) staining. (D) Ctrl HT22 and P301S-Tau HT22 cells were cultured in medium with specified glucose concentrations for up to 48 hours, with the treatment of Ferr-1 (10 µM), z-VAD (20 µM), CQ (25 µM) and DTT (2 mM) for the indicated times. Cell death was assessed using an SG positivity assay. (E) Representative bright field images of cells cultured for 48 hours in medium containing 1.5 mM or 0.7 mM glucose, with or without Nec-1s or GSK-872. Scale bar is 20 µm. (F-G) Immunoblotting of necroptotic markers in Ctrl HT22 and P301S-Tau HT22 cells exposed to low glucose exposure (LGE, 0.7 mM glucose in culture medium) for the indicated time periods. Nec-1s was used to inhibit RIPK1 kinase activity. (H) Representative images of primary cortical neurons isolated from WT and PS19 mice exposed to culture medium containing 1.5 mM or 0.7 mM glucose for 36 hours, with or without Nec-1s. Immunofluorescence staining for MAP2 (green) and AT8 (magenta) is shown. Scale bar is 50 µm. (I) Quantifications of MAP2-positive neuron counts and AT8 staining intensity as shown in (H). (J) Immunoblotting of necroptotic markers in primary cortical neurons isolated from WT and PS19 mice with LGE treatment. Quantified data are presented as mean ± SEM. Two-way ANOVA followed by Dunnett’s multiple comparisons test was used (D) and unpaired two-tailed Student’s t test was used (I). *p < 0.05, **p < 0.01, n.s = not significant.

In tauopathies such as AD, p-Tau accumulation and glucose hypometabolism occur in sequence in the human brain before neuronal loss and brain atrophy^1,2,23^. To explore the possible role of glucose hypometabolism in the death of p-Tau-bearing neurons, we treated P301S-Tau HT22 cells with a culture medium containing a glucose concentration below 1.5 mM. We observed that reducing glucose levels in the culture medium induced a concentration-dependent increase in the death of P301S-Tau HT22 cells, but not in control cells (Figure 1A). To determine whether the glucose hypometabolism-induced cell death observed in P301S HT22 cells is due to the high expression of Tau protein rather than p-Tau, we generated HT22 cells expressing wild-type human Tau protein (WT-Tau HT22) (Figure S1A). WT-Tau HT22 cells express Tau protein at levels similar to P301S-Tau HT22 cells, but show significantly reduced levels of p-Tau. Glucose hypometabolism did not induce cell death in WT-Tau HT22 cells (Figure S1B), suggesting that p-Tau, rather than Tau protein alone, mediates neuronal death under low-glucose conditions.

We next investigated the type of cell death in neurons harboring p-Tau triggered by glucose hypometabolism, utilizing well-established cell death inhibitors. Z-VAD-fmk (Z-VAD) is a cell-permeable pan-caspase inhibitor that effectively blocks apoptosis and pyroptosis^24^. Ferrostatin-1 (Fer-1) inhibits ferroptosis by preventing oxidative lipid damage^25^. Chloroquine (CQ) is an autophagy inhibitor that prevents neuronal cells from oxidative stress^26^. The reducing agent dithiothreitol (DTT) relieves disulfide stress, thereby mitigating disulfidpotosis^27^. Nec-1s^28^ and GSK-872^29^ are potent inhibitors of RIPK1 kinase and RIPK3 kinase activities, respectively, preventing necroptosis. Among these inhibitors examined, only Nec-1s and GSK-872 effectively prevented cell death in P301S-Tau HT22 cells cultured in 1 mM or 0.7 mM glucose (Figure 1B to 1E). Furthermore, treatment with Nec-1s or GSK-872 provided strong protection to P301S-Tau HT22 cells cultured in 0.5 mM glucose, while Ferr-1, Z-VAD, and CQ offered only modest protection, and DTT exacerbated cell death (Figure 1B to 1D). Together, these findings indicate that glucose hypometabolism primarily induces necroptosis in neurons with p-Tau, while other forms of programmed cell death, including apoptosis, pyroptosis, ferroptosis, or oxidative stress, are probably activated when the extracellular glucose concentration falls to an extremely low level.

We further observed that P301S-Tau HT22 cells cultured in medium with low glucose (low glucose exposure, LGE, 0.7 mM glucose in the culture medium) promoted the appearance of necroptosis hallmarks, including p-S166 RIPK1 (pRIPK1), p-T231/232 RIPK3 (pRIPK3) and p-S345 MLKL (pMLKL)^30,31^ (Figure 1F). LGE treatment also led to increased Tau phosphorylation in P301S-Tau HT22 cells, as revealed by AT8 staining (Figure 1F), which is consistent with previous reports^32,33^. In addition to enhanced phosphorylation, LGE treatment was also accompanied by increased Tau aggregation (Figure S1B). In line with its protective effect against cell death in LGE treated P301S-Tau HT22 cells, the RIPK1 kinase inhibitor, Nec-1s, significantly reduced the activation of RIPK3 and MLKL (Figure 1G). Notably, apoptotic and pyroptotic markers, such as the cleavage of caspase-3 and GSDME, were not detected in these cells (Figure S1C). Moreover, LGE did not trigger the activation of the necroptotic markers RIPK3 and MLKL in WT-Tau HT22 cells (Figure S1E), and LGE induced AMPK signaling was comparable among Ctrl, WT-Tau and P301S-Tau HT22 groups (Figure S1F). Adalimumab, a monoclonal anti-TNF antibody widely used to treat TNF associated diseases^34^, failed to rescue LGE induced necroptosis while effectively inhibited TNF induced necroptosis in P301S-Tau HT22 cells (Figure S1G and S1H). These findings suggest that necroptosis driven by glucose hypometabolism in combination with p-Tau occurs independently of the TNF/TNFR1 signaling pathway.

Next, we examined neuronal death induced by glucose hypometabolism in primary cortical neurons isolated from wild-type mice (WT mice) and P301S mutant human Tau transgenic mice (PS19 mice). The isolated neurons were cultured in Neurobasal medium containing a glucose concentration of 25 mM for 10 days in vitro (DIV), followed by exposure to LGE for 36 hours. Neuronal counts were assessed using MAP2 staining, while p-Tau levels were detected by AT8 staining. We found that LGE induced neuronal loss in neurons derived from PS19 mice, but not in those from WT mice, along with elevated p-Tau levels and activation of necroptotic markers (Figure 1H to 1J). In low-glucose conditions, inhibiting RIPK1 kinase activity with Nec-1s, but not other cell death inhibitors including Fer-1, z-VAD, CQ or DTT, effectively prevented neuronal loss in PS19-derived primary neurons without altering p-Tau levels (Figure 1H and 1I, Figure S1I and S1J). Collectively, our data indicate that RIPK1 kinase dependent necroptosis mediates neuronal loss trigged by the synergistic effects of glucose hypometabolism and p-Tau accumulation.

### A20 downregulation is indispensable for neuronal necroptosis triggered by the synergistic effects of p-Tau and low-glucose conditions

Since p-Tau expressing primary neurons and HT22 cells cultured in low glucose containing medium is sufficient to trigger RIPK1 activation and subsequent necroptosis without TNF stimulation, we next explored the mechanism by which low glucose conditions might sensitize the activation of RIPK1 in neurons harboring p-Tau. A20, also known as TNFAIP3, is a key negative regulator of necroptosis^18,19^. Previously, we showed that reduced plasma membraned GLUT1 can impair glucose uptake, leading to decreased acetyl-CoA levels, which in turn reduces lysine acetylation and promotes lysosomal degradation of A20^35,36^. Therefore, we hypothesized that glucose hypometabolism might likewise result in A20 downregulation. Indeed, we observed a time-dependent decrease in A20 levels following LGE treatment in both Ctrl and P301S-Tau HT22 cells, as well as in primary cultured neurons (Figure 2A, Figure S2A). To assess whether A20 downregulation plays a critical role in glucose hypometabolism-induced necroptosis, we transfected P301S HT22 cells with a lentivirus expressing A20 or an empty vector. We found that A20 overexpression significantly protected against necroptosis induced by LGE treatment in neurons harboring p-Tau (Figure 2B and 2C).

**Figure 2.**
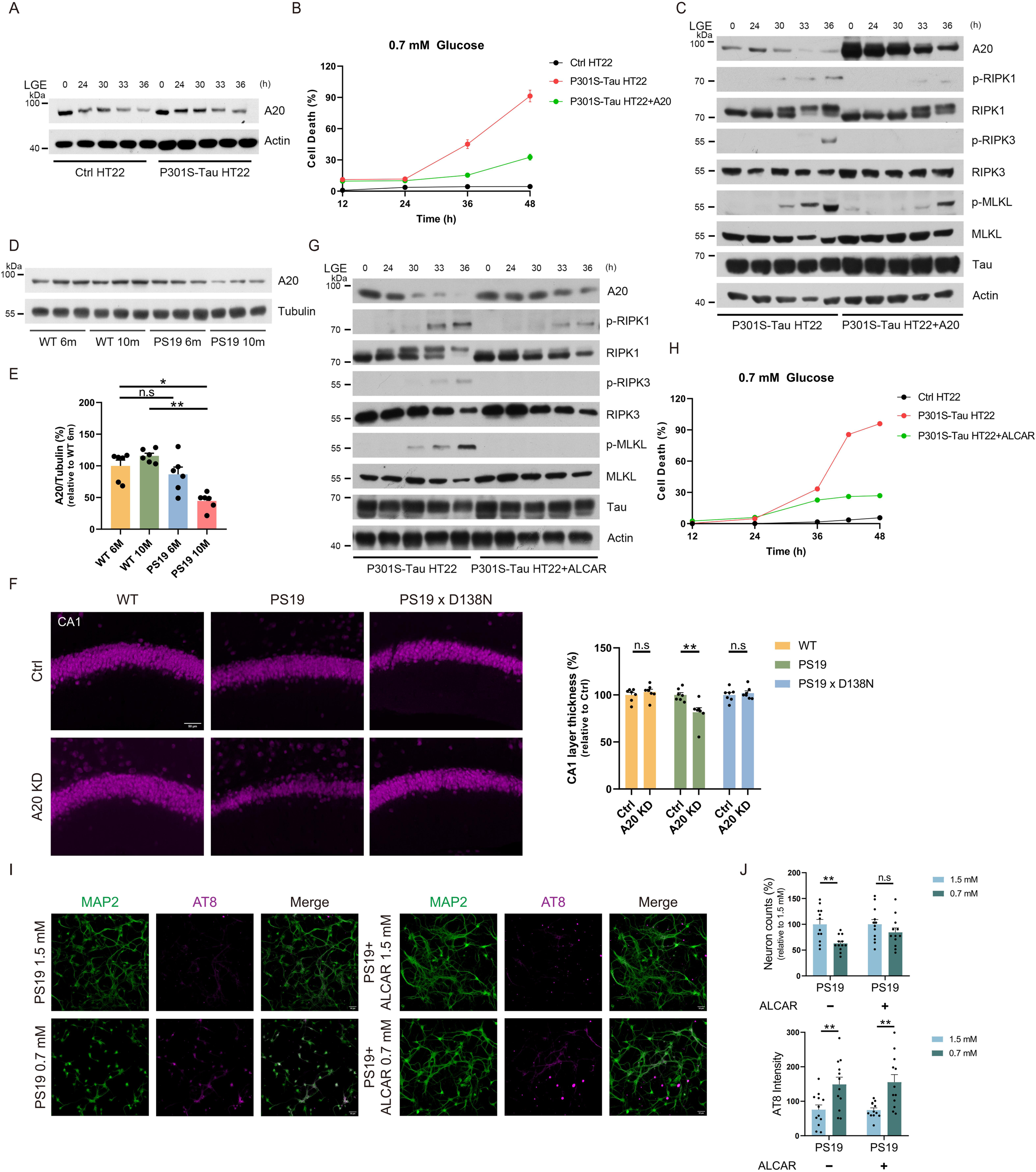
Downregulation of A20 is indispensable for necroptosis triggered by the synergistic effects of p-Tau and glucose hypometabolism. (A) Immunoblotting of A20 levels upon low glucose exposure (LGE, 0.7 mM glucose in culture medium) treatment in Ctrl and P301S-Tau HT22 cells. (B-C) Ctrl HT22 cells were transfected with lentivirus expressing an empty vector and P301S-Tau HT22 cells transfected with lentivirus expressing A20 or an empty vector were cultured in medium containing 0.7 mM glucose. Cell death was assessed using an SYTOX Green (SG) positivity assay (B). Protein levels of the indicated targets were determined by immunoblotting (C). (D-E) Immunoblotting (D) and quantification (E) of A20 expression levels in hippocampal lysates from WT or PS19 mice. n = 6 mice/group. (F) Immunofluorescence analysis of pyramidal cell layer thickness in the CA1 region using NeuN staining in WT, PS19 and PS19xD138N mice, three weeks after injection with A20-shRNA or Ctrl AAV. PS19xD138N mice are generated by crossing PS19 tau transgenic mice with RIPK1 kinase–dead D138N knock-in mice. n = 7 mice per group. Scale bar is 50 µm. (G-H) P301S-Tau HT22 cells cultured in medium containing 0.7 mM glucose with or without acetyl-L-carnitine (ALCAR, 20 mM) for the indicated durations. Cell death was assessed by SG positivity assay (H). The levels of indicated proteins were determined by immunoblotting (G). (I-J) Representative images and quantification of primary cortical neurons isolated from WT and PS19 mice exposed to culture medium containing 1.5 mM or 0.7 mM glucose for 36 hours, with or without ALCAR (20 mM). Immunofluorescence staining for MAP2 (green) and AT8 (magenta) is shown. Quantification of MAP2-positive neuron numbers and AT8 intensity is presented in (J). Scale bar is 50 µm. Quantified data are presented as mean ± SEM. Kruskal–Wallis test followed by Dunn’s multiple comparisons test was used (E) and two-tailed unpaired Student’s t test was used (F and J). *p < 0.05, **p < 0.01, n.s = not significant.

FDG-PET imaging revealed glucose hypometabolism in the hippocampus of PS19 mice, together with a time-dependent decline in neuronal glucose transporter GLUT3 expression and hippocampal volume in these mice^37–39^. We found that A20 expression levels were decreased in the hippocampus of PS19 mice at 10 months but remained unchanged at the age of 6 months (Figure 2D and 2E). The downregulation of neuronal A20 in aged PS19 mice was further confirmed by flow cytometry (Figure S2B and S2C). A20 is a key regulator of necroptosis, and previous studies have shown that pharmacological inhibition of necroptosis prevents neuronal degeneration in Tau transgenic mice^5,40^. Consistent with this, we further demonstrate that neuronal MLKL deficiency achieved via AAV delivery under a neuronal promoter effectively prevents neuronal loss and cognitive dysfunction in PS19 mice (Figure S2D to S2L).

To further explore whether A20 deficiency could contribute to neuronal necroptosis in PS19 mice, we introduced A20-shRNA or Ctrl-shRNA via AAV into the hippocampus of 6-month-old PS19 mice and WT littermates. After three weeks, analysis of the dissected hippocampus confirmed decreased A20 expression via immunoblotting (Figure S3A). Knockdown of A20 resulted in a significantly thinner pyramidal cell layer in the CA1 region of PS19 mice, accompanied by elevated neuronal necroptotic signaling, as indicated by increased pRIPK1 and pMLKL, an effect that was not observed in WT mice (Figure 2F, Figure S3B to S3E). Importantly, genetic inactivation of RIPK1 kinase through the RIPK1 D138N mutation in PS19 mice effectively prevented neuronal loss and neuronal necroptosis caused by A20 deficiency (Figure 2F, Figure S3B to S3E). These findings collectively highlight a crucial role for A20 in regulating p-Tau-mediated neuronal necroptosis in vivo and suggest that the reduction of A20 might provide a hit for promoting necroptosis in neurons that express p-Tau.

Acetyl-L-carnitine (ALCAR), a dietary supplement previously shown to improve cognitive function in AD patients^41,42^, is a metabolic precursor of acetyl-CoA, a key donor of acetyl group in cellular biochemical reactions. We previously showed that ALCAR treatment could increase lysine acetylation of A20 and therefore prevent its lysosomal degradation caused by impaired glucose uptake^35,36^. Consistent with previous findings, here we found that treatment with ALCAR effectively, though not completely, reduced necroptosis in P301S-Tau HT22 cells and in primary neurons isolated from PS19 mice under low-glucose conditions accompanied by increased A20 expression (Figure 2G to 2J, Figure S3F). These data indicate that ALCAR treatment significantly attenuates necroptosis induced by the combined effects of glucose hypometabolism and p-Tau, while residual necroptotic signaling below the detection threshold of Western blot analysis or engagement of non-necroptotic cell death pathways may still contribute to a low level of cell death under these conditions. Moreover, A20 knockdown in P301S-Tau HT22 cells, but not Ctrl HT22 cells, markedly enhanced necroptosis trigged by glucose hypometabolism, which was no longer rescued by ALCAR treatment, whereas Nec-1s and GSK-872 remained effective (Figure S3G to S3I). These findings suggest that the protective effects of ALCAR against necroptosis induced by the combined effects of p-Tau and glucose hypometabolism are dependent on A20.

Taken together, these results demonstrate that A20 deficiency provides the mechanism downstream of glucose hypometabolism to trigger necroptosis in neurons with p-Tau. Furthermore, genetically or pharmacologically enhancing A20 expression offers an effective strategy to prevent necroptosis in p-Tau-expressing neurons under glucose hypometabolism condition.

### ALCAR treatment restores A20 expression and mitigates neurodegeneration in PS19 mice

To assess whether ALCAR as a dietary supplement could reverse A20 downregulation in aged PS19 mice, we administered ALCAR in drinking water beginning at 6 months of age^38^. By 10-11 months of age, ALCAR supplement effectively restored A20 expression in the hippocampus of PS19 mice (Figure 3A). Immunochemistry (IHC) confirmed reduced A20 expression in the pyramidal neurons of PS19 mice, which was restored following ALCAR supplementation (Figure 3B). Interestingly, we observed elevated A20 expression in microglia of PS19 mice (Figure S4A), consistent with prior reports that activated microglia exhibit increased glucose uptake and upregulate A20 expression^43,44^. This activated microglial response may be dampened by ALCAR treatment, leading to reduced A20 expression in microglia (Figure S4B). Additionally, ALCAR treatment attenuated microglia activation and concurrently reduced A20 expression in microglia (Figure S3B).

**Figure 3.**
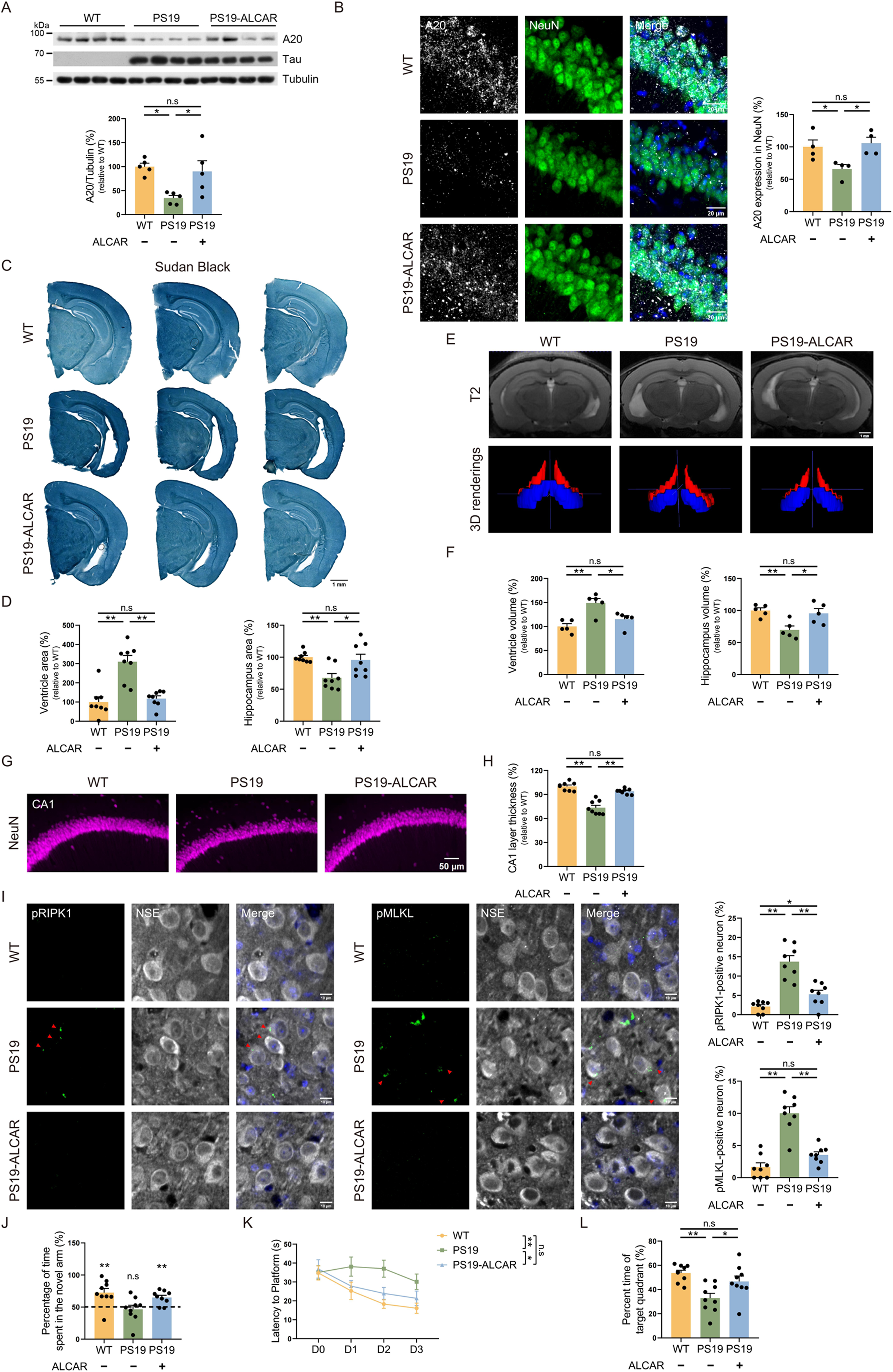
Acetyl-L-carnitine treatment restores A20 expression levels and rescues neurodegeneration in PS19 mice. (A) Immunoblotting and quantification of A20 expression levels in hippocampal lysates from ALCAR treated PS19 mice. n = 5 mice/group. (B) Representative images and quantification of A20 immunostaining in CA1 neurons of ALCAR treated PS19 mice. A20 (gray), NeuN (green), Dapi (blue) staining are shown. n = 4 mice/group. Scale bar is 20 µm. (C-D) Representative images of serial mouse brain sections stained with Sudan black (C), and quantification of lateral ventricle and hippocampus size in WT and PS19 mice treated with or without ALCAR (D). n = 8 mice/group. Scale bar is 1 mm. (E-F) Representative images (E) and quantification (F) of T2 MRI scans from ALCAR treated PS19 mice. 3D renderings of the hippocampus (blue) and lateral ventricle (red) are shown in the lower panels of (E). n = 5 mice/group. Scale bar is 1mm. (G-H) Immunofluorescence analysis of pyramidal cell layer thickness via NeuN staining in the CA1 region of ALCAR treated PS19 mice. n = 8 mice/group. Scale bar is 50 µm. (I) Representative images and quantification of pRIPK1 and pMLKL immunostaining in the CA1 region of ALCAR treated or untreated PS19 mice. Colocalization of pRIPK1 or pMLKL (green, indicated by red arrowheads) with NSE (gray) and Dapi (blue) are shown. n = 8 mice/group. Scale bar is 10 µm. (J-L) Cognitive behavior analysis of ALCAR-treated PS19 mice. (J) Novel arm preference, quantified as the percentage of time spent in the novel arm, during the T-maze test. (K) Escape latency during training trials and (L) target quadrant preference, quantified as the percentage of time spent in the target quadrant, during the probe test of Morris water maze. n = 9 mice/group. Quantified data are presented as mean ± SEM. One-way ANOVA followed by Tukey’s multiple comparisons test was used (A, B, D, F, H, I and L); One sample t test against 50% was used (J); two-way ANOVA followed by Tukey’s multiple comparisons test was used (K). *p < 0.05, **p < 0.01, n.s = not significant.

Importantly, ALCAR administration significantly alleviated hippocampal atrophy and decreased lateral ventricle enlargement in PS19 mice, as quantified from brain sections stained with Sudan Black (Figure 3C and 3D). To further assess brain atrophy in living PS19 mice, we conducted magnetic resonance imaging (MRI). Using three-dimensional rendering of T2-weighted images, we quantified the volumes of lateral ventricle and hippocampus. Volumetric analysis revealed that ALCAR treatment significantly reversed brain atrophy in PS19 mice, marked by reduced lateral ventricle size and increased hippocampal volume (Figure 3E and 3F). Additionally, ALCAR restored the thickness of the pyramidal cell layer in the CA1 region as revealed by immunofluorescence (Figure 3G and 3H). These findings suggest that brain atrophy and neurodegeneration of PS19 mice could be rescued by ALCAR treatment.

ALCAR supplementation successfully prevented neuronal necroptosis in PS19 mice at 10-11 months of age, as indicated by the reduction in the levels of pRIPK1 and pMLKL (Figure 3I). Additionally, ALCAR treatment effectively mitigated spatial memory deficits in PS19 mice, as assessed by the novel T maze test and water maze test (Figure 3J to 3L). Taken together, our findings suggest that restoring neuronal A20 expression via ALCAR as a dietary supplement may represent a promising therapeutic strategy to protect against necroptosis-driven cognitive dysfunction in PS19 mice.

### Assessing A20 expression and RIPK1 activation across hippocampal subregions exhibiting distinct glucose uptake profiles in AD patients

Brain glucose metabolism is highly sensitive to postmortem delay, making it challenging to compare glucose metabolism–regulated proteins, such as A20, between control and AD patient samples due to variability in tissue collection times. To address this limitation, we examined the relationship between glucose uptake, A20 expression, and RIPK1 activation by analyzing subregional differences within the hippocampus of individual AD patients, as this within-patient subregional analysis circumvents inter-sample variability in postmortem delay.

In three AD cases that underwent FDG-PET scanning during lifetime and later postmortem examination, we observed that the subiculum had markedly higher glucose uptake than the CA regions (Figure 4A). This result is consistent with a large-scale analysis of 846 participants, including both AD patients and healthy controls, which also reported significantly higher glucose uptake in the subiculum than in the CA regions (summarized in Figure 4B)^45^. Notably, although neurofibrillary tangles (NFTs) are present in both regions, significant loss of MAP2-positive neurons was detected only in the CA regions of intermediate-stage AD cases (summarized in Figure 4B)^46^. Aligned with their relatively lower glucose uptake and increased susceptibility to AD pathogenesis, the CA regions exhibited diminished A20 expression in neurites and an increased number of neurons positive for pRIPK1 compared to the subiculum (Figure 4A and 4C). The overall intensity of A20 and pRIPK1 immunostaining in these two hippocampal subregions followed a similar trend but did not reach statistical significance in this small sample (Figure S5A).

**Figure 4.**
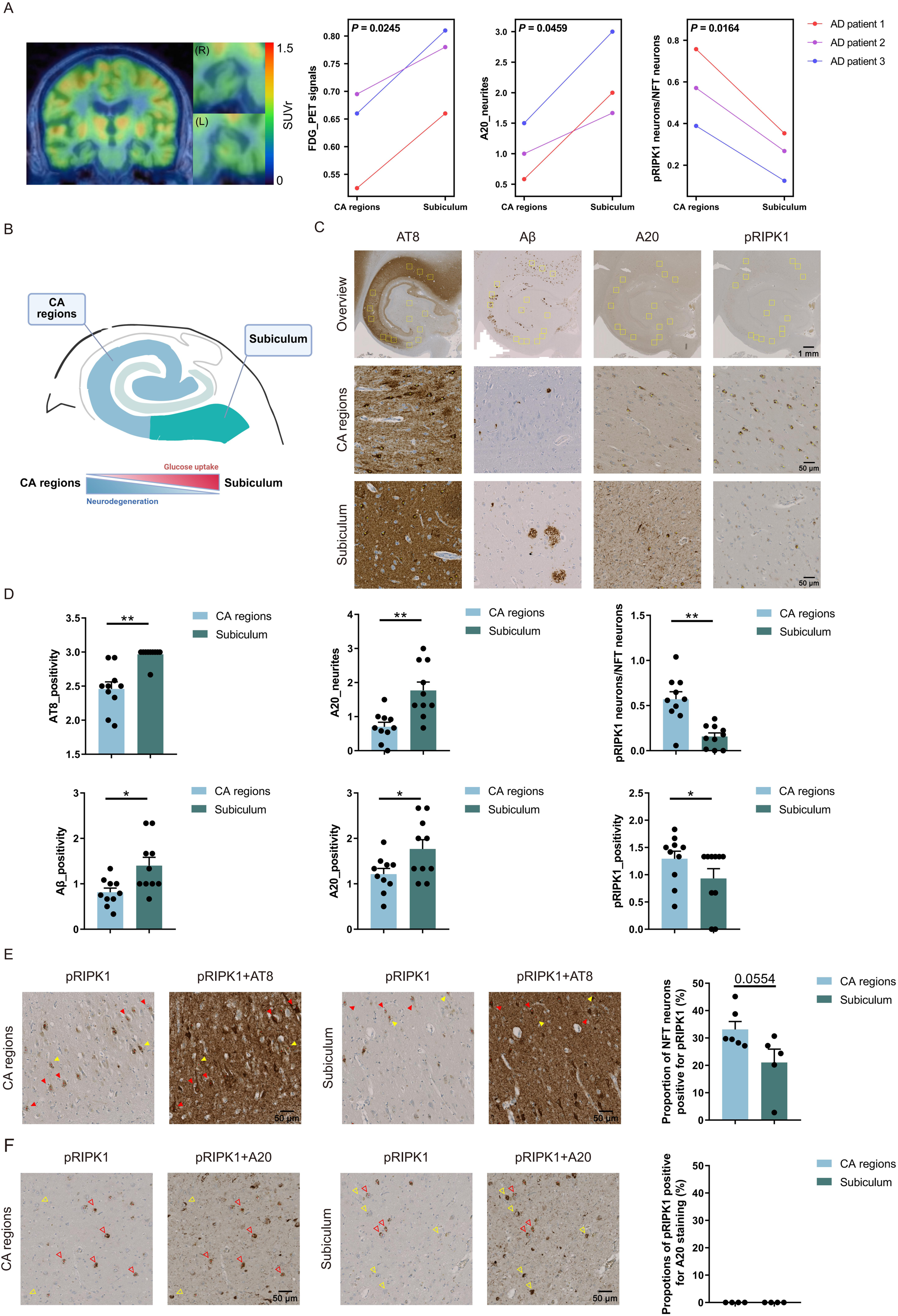
Assessment of A20 and pRIPK1 expression in hippocampal subregions of AD patients. (A) Representative images of FDG-PET imaging in one AD patient show coronal overview and hippocampus zooms upon a high-resolution MRI (T1-weighted, isotropic 1 mm voxels) after scaling to cerebellar grey matter intensity (left). FDG_PET signals refer to standardized uptake value ratios (SUVR). Quantification of FDG-PET signals, A20 and pRIPK1 immunostaining in hippocampal CA regions and the subiculum of three AD patients that underwent FDG-PET scanning during lifetime (right). (B) Schematic diagram illustrating differences in glucose uptake and neurodegeneration between subiculum and CA regions of AD patients based on published reports. (C) Representative images show AT8, Aβ, A20 and pRIPK1 immunostaining in hippocampal overview (top), CA regions (middle) and the subiculum (bottom) of AD patients. The yellow boxes in the overview image indicate the 500 µm x 500 µm regions selected for quantification. Scale bars are 1mm (overview) and 50 µm (magnified view). (D) Quantification of AT8, Aβ, A20 and pRIPK1 immunostaining in hippocampal CA regions and the subiculum of AD patients. For pRIPK1 neurons/NFT neurons, the proportion was quantified by counting each population separately on their respective sections, followed by calculation of the ratio. n = 10 patients. (E) Representative images and quantification of sequential staining for pRIPK1 and AT8 at the same brain section. pRIPK1 positive neurons are labeled with red dots. Exemplary, red solid arrowheads point to double positive neurons and yellow solid arrowheads point to pRIPK1 positive and NFT negative neurons. The proportion of NFT neurons positive for pRIPK1 was calculated by double positive neurons/NFT neurons in the same region of the overstaining. (F) Representative images and quantification of Sequential staining for pRIPK1 and A20 at the same brain section. pRIPK1 positive neurons are labeled with red dots. Exemplary, red empty arrowheads point to pRIPK1 positive neurons and yellow empty arrowheads point to A20 positive neurons. There was no indication of pRIPK1_A20 double positivity because neurons positive for pRIPK1 did not show an increase in signal with sequential A20 staining. Quantified data are presented as mean ± SEM. A paired two-tailed t-test was used (A); Mann-Whitney U test was used (D) and unpaired Student’s t test was used (E). *p < 0.05, **p < 0.01, n.s = not significant.

To expand upon these findings and further test the association between glucose uptake, A20 expression, and RIPK1 activation, we analyzed seven additional AD patient hippocampi. While these patients did not undergo FDG-PET scanning, we speculate that the subiculum generally has higher glucose uptake than the CA regions based on a large-cohort study^45^. Hematoxylin and eosin (H&E) staining showed that neuronal density was comparable between the subiculum and CA areas in these samples (Figure S5B). The results demonstrated that AT8-positive p-Tau, 4G8-positive Aβ pathology and neuronal NFT burden were more pronounced in the subiculum than in the CA regions (Figure 4D, Figure S5C). Despite this, and consistent with the subiculum’s higher expected glucose uptake, the subiculum exhibited elevated A20 expression, both within neurites and in overall staining, relative to the CA regions (Figure 4D). In contrast, the CA regions demonstrated lower A20 levels and markedly higher pRIPK1 expression in neurons as well as in general staining. This pattern in the CA regions aligns well with their presumed reduced glucose uptake and documented susceptibility to neurodegeneration in AD^46^ (summarized in Figure 4B).

To further explore the potential association among Tau phosphorylation, RIPK1 activation, and A20 expression in AD, we performed sequential immunostaining for pRIPK1_AT8 and pRIPK1_A20 on the same hippocampal sections from AD patients. We observed that a substantial fraction of AT8-positive NFT-bearing neurons exhibited RIPK1 activation, accounting on average for 33% of NFT-bearing neurons in the CA regions and 21% in the subiculum (Figure 4E). In contrast, pRIPK1-positive neurons were consistently negative for A20 expression (Figure 4F). These findings support a cellular-level association between Tau phosphorylation, A20 downregulation, and RIPK1 activation in AD patient brains.

Taken together, while these human data alone do not establish causality, they indicate a correlative relationship between regional glucose uptake, A20 expression, and RIPK1 activation in p-Tau affected hippocampal subregions of AD patients, in line with mechanistic insights obtained from the cell and mouse models used in this study.

### RIPK1 interacts with p-Tau via an evolutionarily conserved charged residue-rich sequence

While A20 downregulation is important in driving glucose hypometabolism–induced necroptosis in neurons harboring p-Tau, A20 deficiency alone is insufficient to trigger neuronal necroptosis in Ctrl cells (Figure 2A, Figure S3I) or neurodegeneration in WT mice (Figure 2F). We thus explored other pathological factors that might be associated with glucose hypometabolism to promote necroptosis. Consistently, we observed that p-Tau levels increased in low glucose conditions (Figure 1F and 1J), a response previously connected to p38 activation^32^. Activated p38 MAP kinase has been shown to directly phosphorylate Tau, and inhibition of p38 MAPK has been explored as a potential therapeutic strategy for AD^47,48^. We thus hypothesized that elevated p-Tau levels downstream of p38 activation might contribute to necroptosis induced by glucose hypometabolism. Supporting this, pharmacological inhibition of p38 MAPK signaling effectively suppressed the LGE-induced increase in p-Tau levels and attenuated necroptosis of P301S-Tau HT22 cells in low-glucose conditions (Figure 5A and 5B). Notably, p38 inhibition did not restore A20 expression.

**Figure 5.**
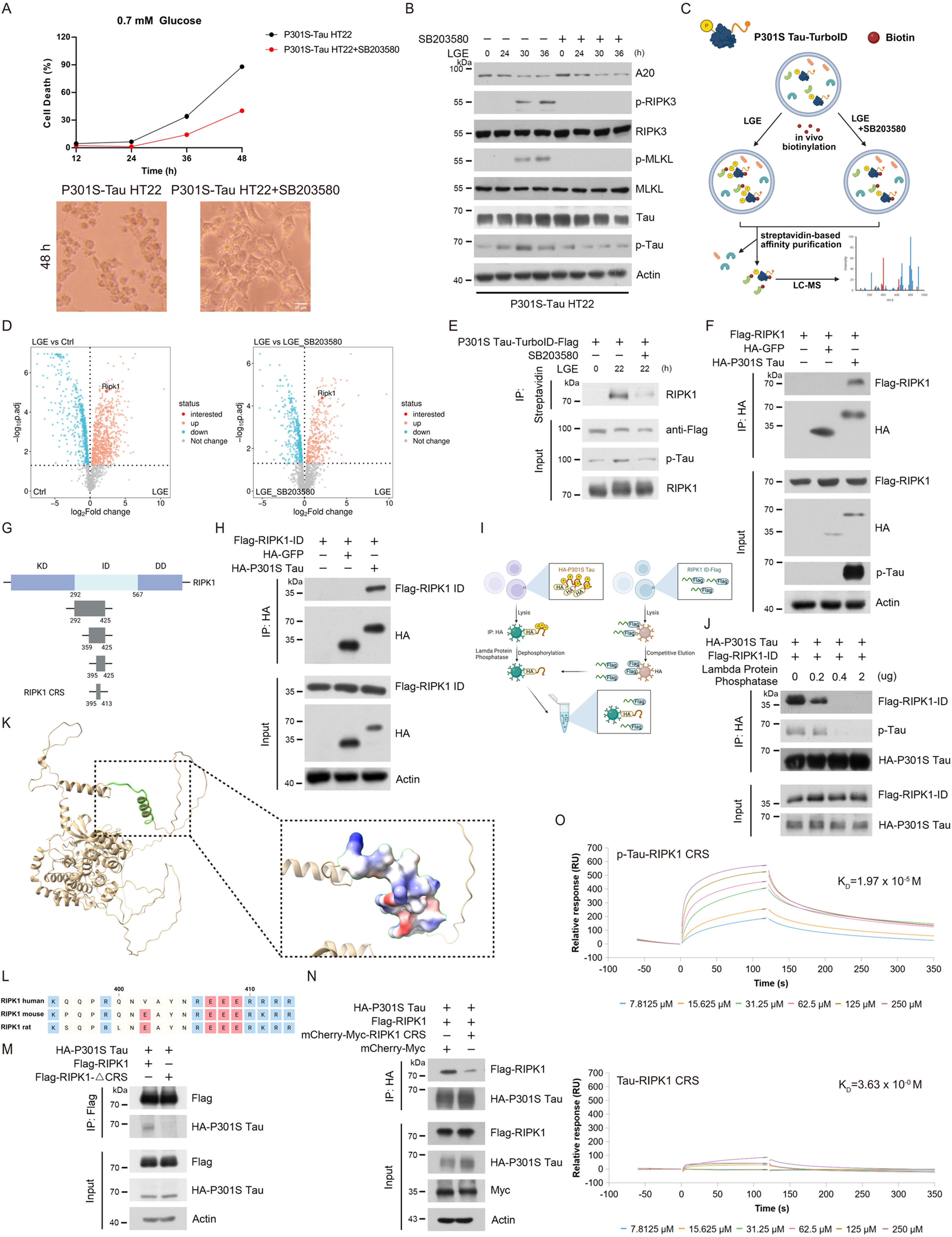
p-Tau interacts with RIPK1 through a charged residue-rich sequence. (A-B) P301S-Tau HT22 cells were cultured in medium containing 0.7 mM glucose with or without the p38 MAPK inhibitor SB203580 (2.5 μM), for the indicated durations. Cell death was assessed by SYTOX Green (SG) staining (A). Representative bright field images of cells cultured for 48 hours in medium containing 0.7mM glucose with or without SB203580. The levels of indicated proteins were determined by immunoblotting (B). Scale bar is 20 µm. (C) Schematic diagram of the proximity labeling workflow for identifying proteins interacting with p-Tau. (D) Volcano plot showing the distribution of biotin-labeled proteins pulled down from P301S Tau-TurboID expressing cells, comparing conditions before and after low glucose exposure (LGE, 0.7 mM glucose in culture medium) treatment. SB203580 was used to dephosphorylate p-Tau during LGE treatment. (E) The interaction between p-Tau and RIPK1 identified by biotin proximity labeling in (D) was confirmed via immunoblotting. (F) HEK 293 cells were transiently transfected with HA-tagged P301S Tau or HA-tagged GFP, together with Flag-tagged RIPK1, for 48 hours. Cell lysates were immunoprecipitated using anti-HA-conjugated beads, and the interaction between HA-P301S Tau and Flag-RIPK1 was detected by immunoblotting. (G) Schematic diagram of RIPK1 truncation constructs. (H) HEK 293 cells were transfected for 48 hours with plasmids encoding HA-tagged P301S Tau or HA-tagged GFP, together with Flag-tagged RIPK1 ID domain. Cell lysates were subjected to immunoprecipitation using anti-HA-conjugated beads. (I-J) HEK 293 cells were transfected with plasmids encoding HA-tagged P301S Tau, and lysates were subjected to immunoprecipitation using anti-HA-conjugated beads. After washing away unbound proteins, the beads were treated with varying concentrations of lambda protein phosphatase for 1 hours. The reaction was quenched with 50 mM EDTA before incubating the beads with lysates containing Flag-tagged RIPK1 ID domain. A schematic illustration of the experimental procedure is shown in (I). Interactions between p-Tau and the RIPK1 ID domain were assessed by immunoblotting (J). (K) A schematic diagram of the RIPK1 protein structure predicted by Alpha-fold is shown with the charge distribution of the RIPK1 CRS domain highlighted. (L) Schematic representation of RIPK1 sequence alignment across different species. (M) HEK 293 cells were transfected with expression plasmids of HA-tagged P301S Tau and Flag-tagged RIPK1 or RIPK1 CRS truncated mutant. The cell lysates were then immunoprecipitated with anti-Flag-conjugated beads. Immunoprecipitates were analyzed by immunoblotting. (N) HEK 293 cells were co-transfected with expression plasmids encoding HA-tagged P301S Tau, Flag-tagged RIPK1, and either mCherry-Myc-RIPK1 CRS or mCherry-Myc control. Cell lysates were subjected to immunoprecipitation using anti-HA-conjugated beads, and immunoprecipitates were analyzed by immunoblotting. (O) Surface plasmon resonance (SPR) binding analysis was performed to quantify the molecular interactions between p-Tau (top) or non-phosphorylated (bottom) Tau and the RIPK1 CRS domain. Recombinant Tau protein was expressed and purified from E. coli, followed by in vitro phosphorylation to generate p-Tau.

To elucidate the mechanisms by which p-Tau accumulation contributes to necroptosis, we employed a proximity-labeling strategy combining Turbo-ID with mass spectrometry (Figure 5C). This unbiased approach enabled the identification of candidate p-Tau-interacting proteins that may mediate glucose hypometabolism induced necroptosis (Figure 5C). Unexpectedly, Turbo-ID labeling revealed a strong interaction between RIPK1 and Tau following LGE treatment (Figure 5D). Moreover, inhibition of p38 MAPK signaling reduced this RIPK1-Tau association under low-glucose conditions, indicating that elevated p-Tau levels are critical for facilitating this interaction (Figure 5D). These mass spectrometry findings were further validated by immunoblotting (Figure 5E). These findings suggest that glucose hypometabolism might promote the Tau-RIPK1 interaction by increasing the phosphorylation levels of Tau.

Next, we co-transfected HEK 293 cells with plasmids encoding the P301S Tau protein and RIPK1. Upon overexpression, P301S Tau exhibited high levels of phosphorylation and was found to co-immunoprecipitated with RIPK1 (Figure 5F). RIPK1 comprises a kinase domain (KD), a death domain (DD), and an intermediate domain (ID) (Figure 5G). To determine which domain of RIPK1 is responsible for its interaction with tau, we generated truncated versions of RIPK1, each lacking one of these domains as indicated. Our results showed that deletion of the ID eliminated the interaction between RIPK1 and P301S Tau (Figure S6A). Furthermore, the ID of RIPK1 alone was sufficient to mediate binding to Tau (Figure 5H). Using in vitro phosphatase-treated P301S Tau, we demonstrated that the RIPK1 ID preferentially co-immunoprecipitated with p-Tau, but not with its unphosphorylated form (Figure 5I and 5J).

To pinpoint a more specific region within the ID of RIPK1 that mediates its interaction with p-Tau, we systematically generated a series of truncated segments of the RIPK1 ID region (Figure S6B). Our findings revealed that a charged residue-rich sequence (CRS), spanning residues K395 to R413 in murine RIPK, was capable of interacting with p-Tau (Figure S6B). This CRS is highly conserved across human, mouse and rat, and AlphaFold predicts it to adopt an alpha-helix structure (Figure 5K and 5L). To further validate the necessity of the RIPK1 CRS for the p-Tau-RIPK1 interaction, we generated a truncated version of RIPK1 lacking the CRS (RIPK1-△CRS). The deletion effectively abolished the interaction between overexpressed P301S Tau and RIPK1 (Figure 5M). Additionally, overexpression of RIPK1 CRS could compete with full-length RIPK1 for p-Tau binding, as evidenced by a decrease in the co-immunoprecipitation between RIPK1 and P301S Tau protein (Figure 5N). These results demonstrate that the RIPK1 CRS is critical for mediating the interaction between p-Tau and RIPK1.

To assess the binding affinity between the RIPK1 CRS and p-Tau, we next employed the surface plasmon resonance (SPR) assay. This analysis included both unphosphorylated Tau isolated from E. coli and its in vitro phosphorylated form (p-Tau) via p38 as previously described^49^(Figure S6C). The RIPK1 CRS exhibited no detectable binding to unphosphorylated Tau, whereas it bound p-Tau with an equilibrium dissociation constant (K_D_) of 19.7 μM (Figure 5O). Similarly, in vitro phosphorylation of Tau using another kinase, MARK2^50^, resulted in comparable binding to RIPK1 CRS (Figure S4D, KD = 37.3 μM). Together, these results underscore the critical role of Tau phosphorylation in enabling its interaction with the RIPK1 CRS.

To further examine the molecular basis of the interaction between RIPK1 CRS and p-Tau, we performed all-atom molecular dynamics (MD) simulations focused on the RIPK1 CRS domain and a triply phosphorylated Tau fragment corresponding to the epitope recognized by the AT8 antibody used in this study to detect p-Tau^51^. MD simulations revealed that p-Tau-F preferentially binds to the RIPK1 CRS compared to its unphosphorylated counterpart (Figure S7A). This enhanced interaction was primarily driven by electrostatic forces with the negatively charged phosphate groups on p-Tau-F engaging positively charged arginine residues at positions 12, 16, and 19 of the RIPK1 CRS (Figure S7B and S7C; Videos S1 and S2). To experimentally test the role of these residues, we substituted Arg410 and Arg413 in full-length RIPK1, corresponding to RIPK1 CRS Arg16 and Arg19, with alanine to neutralize their positive charge. As predicted by the MD simulations, these mutations significantly reduced the interaction between RIPK1 and p-Tau, whereas alanine substitutions at adjacent arginine residues (RIPK1 R399A and R412A) had little effect (Figure S7D). Together, these findings suggest that the negatively charged phosphate groups of p-Tau enhance its binding to RIPK1 through electrostatic interactions with specific positively charged residues within the RIPK1 CRS.

### p-Tau condensates serve as a platform for RIPK1 recruitment

Necroptosis is initiated by the assembly of a signaling complex, such as TNF/TNFR1/TRADD, where RIPK1 is recruited via its DD and undergoes dimerization or oligomerization^15,17^. Oligomerized RIPK1, which is then activated through trans-autophosphorylation, subsequently binds to RIPK3 via RHIM domain interactions to form the necrosome, ultimately triggering MLKL activation and membrane rupture^52,53^.

Previous studies have demonstrated that p-Tau readily forms liquid like droplet in vitro, which can further mature into amyloid fibrils^54,55^. Since RIPK1 activation is promoted by its dimerization or oligomerization through trans-autophosphorylation^56^, we examined whether p-Tau condensates are capable of recruiting and potentially activating RIPK1. To this end, we purified full-length human Tau from *E. coli* and generated its phosphorylated form via in vitro phosphorylation^50^. Consistent with prior reports^54,55^, both p-Tau and Tau formed condensates in solution (Figure 6A, labeled with Alexa Fluor 568, AF-568). Upon addition of GFP tagged RIPK1 CRS, we observed selective recruitment and enrichment of RIPK1 CRS within p-Tau condensates, whereas those formed by unphosphorylated Tau did not show such effect. These findings suggest a direct, and phosphorylation-dependent interaction between Tau protein and RIPK1 via the CRS domain.

**Figure 6.**
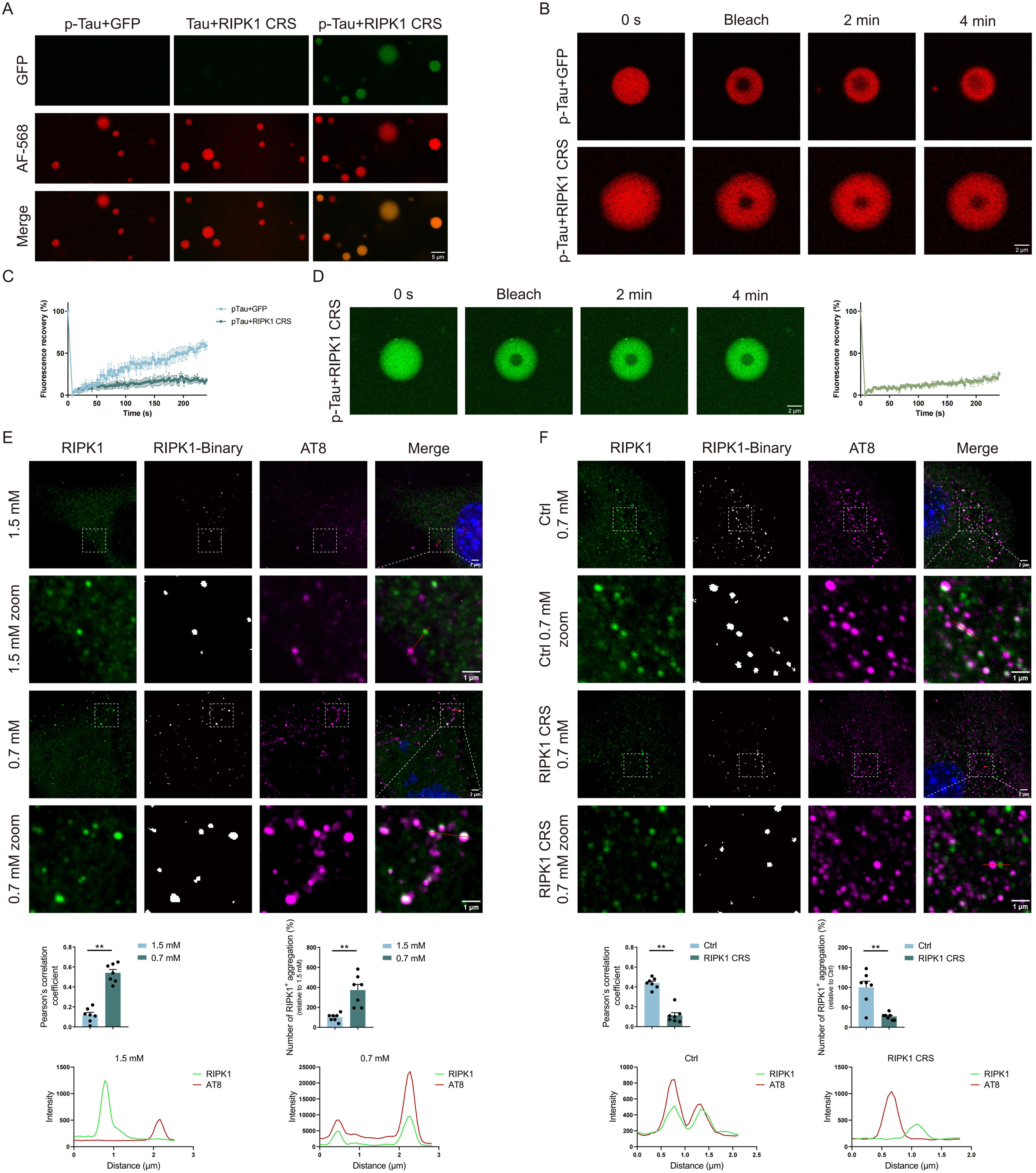
p-Tau condensates recruit RIPK1. (A) Fluorescence images of LLPS formed by Tau or p-Tau with GFP-tagged RIPK1 CRS. Alexa Fluor 568-labeled Tau or p-Tau was mixed with GFP or GFP-RIPK1 CRS in 50 mM Tris (pH 7.5) containing 5% PEG to induce phase separation. Under these conditions, both Tau and p-Tau formed condensates; however, only p-Tau efficiently recruited GFP–RIPK1 CRS, but not GFP into droplets, whereas Tau failed to recruit GFP–RIPK1 CRS. Shown are representative images of the GFP channel (top), AF568 channel (middle), and merged images (bottom). Scale bar is 5 µm. (B) FRAP analysis of p-Tau droplets in the presence or absence of RIPK1 CRS (AF568 channel). p-Tau+GFP droplets (top) exhibited fluorescence recovery after photobleaching, indicating liquid-like behavior. In contrast, p-Tau+RIPK1 CRS droplets (bottom) showed markedly slower recovery, demonstrating reduced fluidity. Images were collected every 2 minutes after bleaching. Scale bar is 2 µm. (C) Quantification of FRAP recovery curves for p-Tau with GFP or GFP-RIPK1 CRS. Fluorescence in the bleached region was normalized to an unbleached control droplet. p-Tau+RIPK1 CRS droplets showed significantly impaired fluorescence recovery relative to p-Tau+GFP, confirming decreased droplet fluidity. (D) FRAP analysis of GFP-RIPK1 CRS within p-Tau condensates. GFP-RIPK1 CRS exhibited minimal fluorescence recovery after bleaching, indicating tight molecular retention and limited exchange within the condensates. Scale bar is 2 µm. (E) P301S-Tau HT22 cells were cultured in medium containing either 1.5 mM or 0.7 mM glucose for 26 hours. Immunostaining displayed the subcellular colocalization of p-Tau (AT8-positive) and RIPK1, and fluorescence intensity histograms of a line scan analyzed by ImageJ are shown below, demonstrating correlated distribution patterns. Colocalization was quantified by calculating Pearson’s correlation coefficients. Similar results were observed in three independent experiments. In addition, RIPK1 aggregation was quantified under these conditions in panel below right. The RIPK1 signal channel was converted into a binary image using a fixed threshold, and this threshold-restricted binary mask was used to highlight RIPK1-positive puncta. Scale bars are 2 µm (overview) and 1 µm (magnified view). (F) P301S-Tau HT22 cells were transduced with lentivirus expressing RIPK1 CRS or empty vector (Ctrl) for 48 hours, followed by treatment with low glucose exposure (LGE, 0.7 mM glucose in culture medium) for 26 hours. Fluorescence intensity histograms of p-Tau and RIPK1 of a line scan analyzed by ImageJ are shown to demonstrate correlated distribution patterns. Colocalization was quantified by calculating Pearson’s correlation coefficients. Similar results were observed in three independent experiments. RIPK1 aggregation was quantified under these conditions in panel below right. Scale bars are 2 µm (overview) and 1 µm (magnified view). Quantified data are presented as mean ± SEM. Unpaired two-tailed Student’s t test was used, **p < 0.01.

We next assessed the dynamic properties of p-Tau condensates in the presence of RIPK1 CRS using fluorescence recovery after photobleaching (FRAP). Condensates containing both p-Tau and RIPK1 CRS exhibited markedly slower fluorescence recovery compared to those formed by p-Tau alone, indicating that RIPK1 CRS incorporation enhanced the solidity of p-Tau condensates (Figure 6B and 6C). Concurrently, RIPK1 CRS displayed limited mobility within p-Tau condensates, indicating that p-Tau not only promotes RIPK1 recruitment but also contributes to its local stabilization within the condensate environment (Figure 6D).

Consistent with these in vitro findings, in P301S HTT2 cells elevated p-Tau levels under low-glucose conditions promoted the recruitment of RIPK1, leading to its aggregation and colocalization with p-Tau (Figure 6E). Notably, overexpression of the RIPK1 CRS domain disrupted this interaction and significantly reduced RIPK1 aggregate formation (Figure 6F). Collectively, these findings suggest that the p-Tau-RIPK1 interaction drives RIPK1 aggregation, which may in turn facilitate its activation.

### The interaction between p-Tau and RIPK1 drives necroptosis triggered by glucose hypometabolism, and disrupting it alleviates neurodegeneration in PS19 mice

To assess whether RIPK1 aggregates formed via interaction with p-Tau contribute to RIPK1 activation and necroptosis, we reconstituted either wild-type RIPK1 or a CRS domain–deficient mutant (RIPK1-ΔCRS) in RIPK1 knockout mouse embryonic fibroblasts (MEFs) stably expressing P301S Tau. Both constructs exhibited comparable expression levels (Figure S8A). While TNF-induced necroptosis and apoptosis responses were similar between the two groups (Figure S8B), exposing cells to low-glucose conditions (0.7 mM) revealed significant differences (Figure 7A). RIPK1-ΔCRS MEFs expressing P301S Tau displayed markedly reduced RIPK1 activation and lower levels of necroptotic marker expression (Figure 7B). This protective effect was also observed in P301S HT22 cells and primary neurons obtained from PS19 mice, where expression of the RIPK1 CRS effectively blocked glucose hypometabolism induced necroptosis (Figure 7C and 7D, Figure S8C and S8D). These findings demonstrate that disrupting the p-Tau-RIPK1 interaction, either by deleting the CRS domain or through competitive inhibition with the CRS domain, effectively suppresses RIPK1 dependent necroptotic signaling triggered by glucose hypometabolism in the context of p-Tau accumulation.

**Figure 7.**
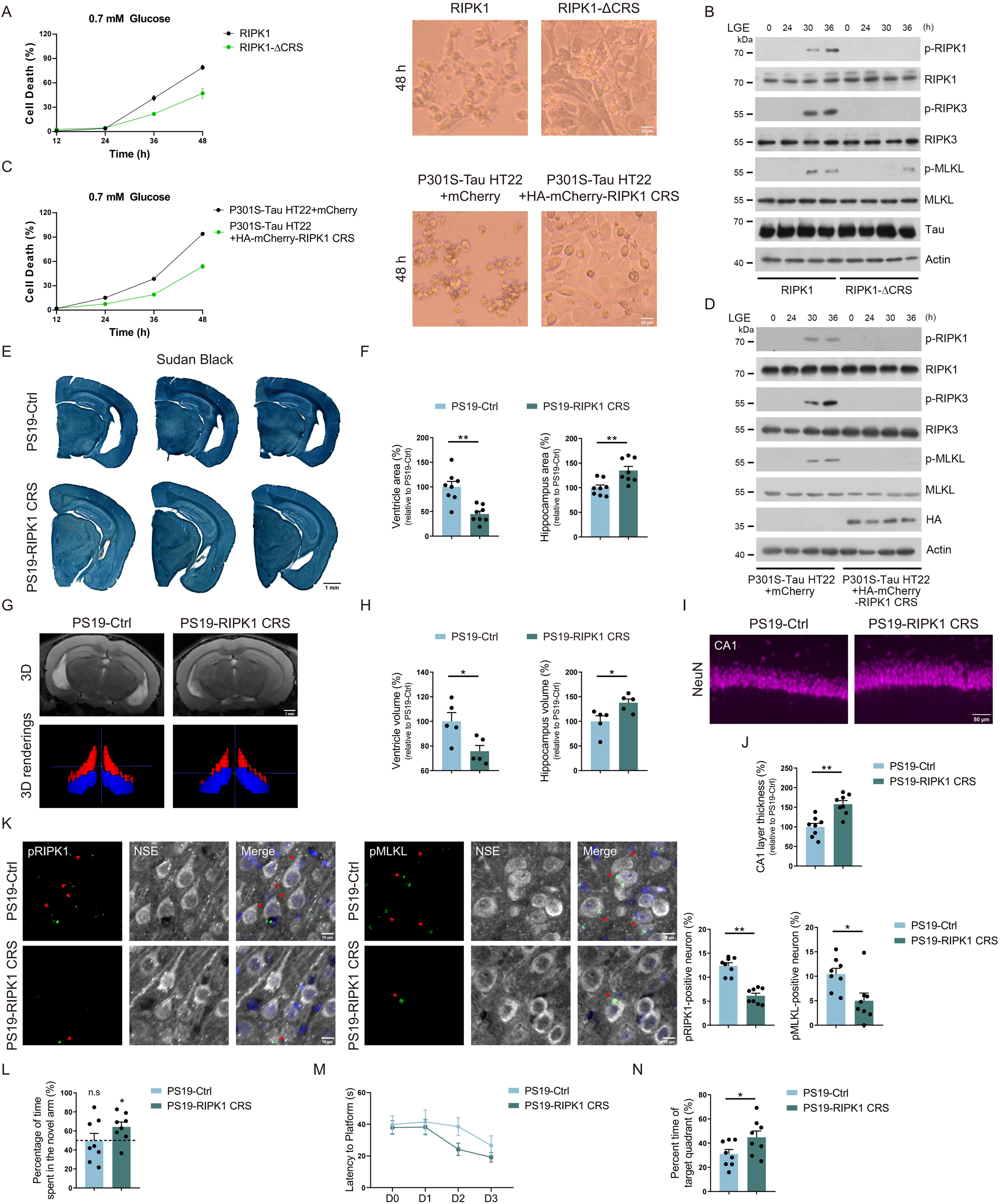
The interaction between p-Tau and RIPK1 drives neurodegeneration in PS19 mice. (A-B) RIPK1 KO MEFs were transfected with lentivirus to stably express either full-length RIPK1 or its CRS truncation mutant (RIPK1-△CRS), along with P301S Tau. Cell death under the treatment of low glucose exposure (LGE, 0.7 mM glucose in culture medium) was assessed by SYTOX Green (SG) staining (A). Representative bright field images of cells cultured for 48 hours in medium containing 0.7 mM glucose. Necroptotic markers exposed to LGE for the indicated time periods were examined by immunoblotting (B). Scale bar is 20 µm. (C-D) P301S-Tau HT22 cells transfected with lentivirus expressing either HA-mCherry-RIPK1 CRS or mCherry were cultured in the medium containing 0.7 mM glucose. Cell death was assessed by SG staining (C). Representative bright field images of cells cultured for 48 hours in medium containing 0.7 mM glucose. The levels of necroptotic markers were assessed by immunoblotting (D). Scale bar is 20 µm. (E-F) Representative images of Sudan black staining in serial brain sections from PS19 mice injected with AAVs encoding either an empty vector (Ctrl) or the RIPK1 CRS domain (E). Quantification of lateral ventricle and hippocampus area is shown (F). n = 8 mice/group. Scale bar, 1 mm. (G-H) Representative images (G) and quantification (H) of T2 MRI scans from PS19 mice injected with AAVs encoding either empty vector (Ctrl) or the RIPK1 CRS domain. 3D renderings of the hippocampus (blue) and lateral ventricle (red) are shown in the lower panels shown in (G). n = 5 mice/group. Scale bar is 1 mm. (I-J) Immunofluorescence analysis of pyramidal cell layer thickness via NeuN staining in the CA1 region of PS19 mice injected with AAVs encoding either empty vector (Ctrl) or the RIPK1 CRS domain. n = 8 mice/group. Scale bar is 50 µm. (K) Representative images and quantification of pRIPK1 and pMLKL immunostaining in the CA1 region of AAV-injected PS19 mice. pRIPK1 or pMLKL (green, indicated by red arrowheads) colocalization with NSE (gray) and Dapi (blue) are shown. n = 8 mice/group. Scale bar is 10 µm. (L-N) Cognitive behavior analysis of AAV injected PS19 mice. (L) Novel arm preference, quantified as the percentage of time spent in the novel arm, during the T-maze test. (M) Escape latency during training trials and (N) target quadrant preference, quantified as the percentage of time spent in the target quadrant, during the probe test of Morris water maze. n = 8 mice/group. Quantified data are presented as mean ± SEM. Unpaired Student’s t test was used (F, H, J, K, N) and One sample t test against 50% was used (L). *p < 0.05, **p < 0.01, n.s = not significant.

Building on our finding that RIPK1 CRS expression inhibits neuronal necroptosis induced by the combined effects of p-Tau pathology and glucose hypometabolism, we investigated the therapeutic potential of RIPK1 CRS in PS19 mice, a model in which both p-Tau accumulation and cerebral hypometabolism progressively emerge. AAV-mediated delivery of RIPK1 CRS into the hippocampal neurons of 6-7-month-old PS19 mice significantly mitigated neurodegeneration by 10-11 months of age (Figure S9A to S9C). CRS-treated PS19 mice showed preserved hippocampal structure, as indicated by increased hippocampal volume and reduced lateral ventricle enlargement, confirmed through both histological (Figure 7E and 7F) and in vivo MRI analyses (Figure 7G and 7H). RIPK1 CRS expression also prevented thinning of the CA1 pyramidal layer (Figure 7I and 7J) and suppressed markers of necroptosis (Figure 7K), verifying inhibition of necroptotic signaling in neurons. Crucially, this intervention rescued cognitive deficits in behavioral tests at 10-11 month old PS19 mice (Figure 7L to 7N, Figure S9G), directly linking the p-Tau-RIPK1 interaction to cognitive decline. Together, these results demonstrate that pathological interaction between p-Tau and RIPK1 drives neuronal necroptosis and neurodegeneration in tauopathies, and that targeting this interaction offers a promising strategy to prevent synergistic neuronal loss caused by p-Tau accumulation and glucose hypometabolism.

## DISCUSSION

Here, we report that glucose hypometabolism sensitizes neurons harboring p-Tau to necroptosis by providing a double-hit involving A20 downregulation and the formation of a p-Tau-RIPK1 signaling hub. A20 acts as a critical checkpoint that restrains necroptosis. However, glucose hypometabolism downregulates A20 expression, thereby lifting this protective brake and permitting necroptotic cell death in neurons harboring p-Tau. Treatment with ALCAR as a dietary supplement enhances A20 lysine acetylation and restores its expression under low-glucose conditions, effectively preventing neuronal loss and cognitive decline in PS19 mice. In addition to A20 downregulation, we show that p-Tau accumulation under low glucose conditions facilitates the recruitment of RIPK1, forming a signaling platform that initiates necroptosis. Moreover, disrupting the interaction between RIPK1 and p-Tau using a RIPK1-derived peptide mitigates necroptosis and neuronal loss in PS19 mice. Together, these findings identify a necroptotic signaling axis initiated by the p-Tau–RIPK1 complex and underscore the contribution of glucose hypometabolism–induced necroptosis to the pathogenesis of tauopathies.

Activation of necroptosis in neurons, evidenced by the phosphorylation of RIPK1, RIPK3, and MLKL, has been observed in p-Tau associated transgenic mouse models, xenografted human neurons, and postmortem brain samples from patients^3–5^. Inhibiting RIPK1 or RIPK3 kinase activity to prevent neuronal necroptosis holds therapeutic potential for reducing neuronal loss, thereby preserving cognitive function in AD or other tauopathies^5,36,40,57,58^. To expand opportunities for targeting this pathway or to further identify the therapeutic window in tauopathies, it is essential to investigate the upstream regulators and triggers of neuronal necroptosis during Tau pathogenesis. A recent report has found that upregulation of the long noncoding RNA MEG3, prompted by amyloid plaques, can induce necroptosis in neurons containing p-Tau^3^; however, the precise mechanism by which MEG3 triggers necroptosis remains unclear. It is worth noting that overexpression of MEG3 has been shown to inhibit glucose metabolism by promoting the degradation of c-Myc, a transcription factor that directly controls the expression of GLUT1 and other glycolytic genes^59,60^. Thus, it is likely that amyloid plaque-induced MEG3 expression could result in glucose hypometabolism and subsequently trigger neuronal necroptosis as a downstream effect of p-Tau accumulation. In our study, we demonstrate that glucose hypometabolism indeed collaborates with p-Tau to induce neuronal necroptosis. Since brain glucose hypometabolism is a common feature and a key driver of neurodegenerative diseases, further research will be needed to identify whether and how MEG3 or other pathological factors differentially expressed in tauopathies contribute to the reduction of brain glucose uptake.

The sequential activation of RIPK1, RIPK3, and MLKL is closely regulated by cell death checkpoints, with A20 being one of them^61^. A20 restricts the ubiquitination of RIPK1 and RIPK3 and thereby protects cells from necroptosis^18,19^. We have previously discovered the role of the glucose-derived metabolite acetyl-CoA in modulating A20 lysine acetylation and regulating its subsequent lysosomal degradation^35,36^. Consistently, here we demonstrate that glucose hypometabolism contributes to A20 deficiency, leading to neuronal necroptosis in tauopathies. Furthermore, restoring A20 levels by providing an alternate source of acetyl-CoA, ALCAR, successfully prevents neuronal loss and brain atrophy associated with p-Tau and glucose hypometabolism. ALCAR treatment for dementia has been evaluated in multiple clinical trials over the past three decades, but inconsistent outcomes have prevented its adoption into routine clinical practice^62–65^. One factor likely contributing to these varied outcomes is probably the lack of a consistent standard for participant selection. Given that our findings demonstrate ALCAR treatment prevents A20 deficiency and neuronal necroptosis caused by glucose hypometabolism, it is reasonable to recommend including FDG-PET scans in ALCAR clinical trials to monitor brain glucose metabolism for patient selection.

The TNFR1 signaling pathway is the most extensively studied mechanism of necroptosis activation^61^. Upon stimulation, trimerized TNFR1 recruits RIPK1 via homotypic DD interactions, leading to the formation of RIPK1-containing puncta that subsequently recruit RIPK3 and activate MLKL^17^. However, our findings demonstrate that TNFR1 signaling is not required for necroptosis triggered by glucose hypometabolism in neurons with p-Tau. Instead, we show that under low-glucose conditions, p-Tau accumulation directly recruits RIPK1, initiating necroptotic signaling. Both in vitro and in vivo experiments reveal that disrupting the interaction between p-Tau and RIPK1 effectively blocks necroptosis induced by the synergistic effects of p-Tau and glucose hypometabolism, thereby mitigating neurodegeneration in PS19 mice. Further research is needed to develop a blood-brain barrier–permeable peptide conjugate to fully harness the therapeutic potential of targeting the p-Tau–RIPK1 interaction.

Necroptosis can also be triggered by TLR3, TLR4, or ZBP1, which directly recruit RIPK3 via their RHIM domain, leading to MLKL activation. In this RHIM-dependent pathway, RIPK1 is generally not required and instead functions as a negative regulator by recruiting caspase-8^66–68^. While we cannot completely exclude the involvement of RIPK1-independent necroptosis in the pathogenesis of tauopathies, both our findings and those of others consistently demonstrate RIPK1 activation in neurons harboring p-Tau, as observed across cultured mouse neurons, xenografted human neurons, multiple transgenic mouse models and postmortem human brain samples^3–5^. Moreover, inhibition of RIPK1 kinase activity has been shown to reduce neuronal necroptosis and prevent cognitive decline associated with neuronal loss^5,40^, supporting a critical role for RIPK1 kinase dependent necroptosis in the progression of tauopathies.

Necroptosis is an inflammatory programmed cell death pathway that leads to the release of damage-associated molecular patterns (DAMPs) and may subsequently trigger a secondary wave of cell death. Notably, activation of the necroptotic pathway has been observed in both p-Tau-positive and p-Tau-negative neurons in postmortem AD patient samples^4^, indicating that there may be at least two distinct mechanisms contributing to the formation of these two types of necroptosis-affected neurons. Given that our data demonstrate glucose hypometabolism-induced necroptosis in tauopathies is dependent on the interaction between p-Tau and RIPK1 rather than external stimuli, we propose that necroptosis in p-Tau-positive neurons could serve as an upstream trigger for p-Tau-negative neurons through the release of DAMPs. Additionally, these DAMPs could activate the NLRP3 inflammasome or upregulate ZBP1 in glial cells or neurons, thereby triggering a secondary release of DAMPs and contributing to the progression of tauopathies^69,70^. On the other hand, neuroinflammation triggered by DAMPs could further disrupt brain glucose metabolism, exacerbating glucose hypometabolism induced necroptosis. Interrupting the detrimental feedback loop of various cell death pathways could open a new avenue for drug development aimed at preventing neuronal loss in tauopathies.

Taken together, our findings address the long-standing question of how glucose hypometabolism and p-Tau synergistically drive neurodegeneration. Beyond providing mechanistic insights, this work proposes a metabolic framework for patient stratification in AD clinical trials. Specifically, we suggest that combined p-Tau and FDG-PET imaging might identify a subset of hypometabolic patients with p-Tau pathology who are most likely to benefit from necroptosis-targeted therapies.

### Limitations of the study

Elevated MEG3 expression has been shown to suppress glucose metabolism by promoting c-Myc degradation and to induce necroptosis in neurons bearing p-Tau. Building on our findings that glucose hypometabolism synergizes with p-Tau to drive neuronal necroptosis, future studies are needed to determine whether MEG3 indeed contributes to impaired neuronal glucose uptake and thereby promotes necroptosis. This line of investigation is particularly important, as the molecular mechanisms underlying brain glucose hypometabolism in AD remain a critical unresolved question in the field. Furthermore, the in vitro phosphorylation of Tau by P38 or MARK2 does not fully recapitulate the phosphorylation patterns of Tau in vivo, representing a limitation in modeling pathological Tau modifications of AD. In addition, the aggregation-prone nature of full-length RIPK1 precluded its purification in vitro, limiting our ability to directly evaluate its binding affinity with p-Tau. Finally, we note that either ALCAR treatment or CRS-mediated deletion of RIPK1 substantially, though not completely, prevented cell death driven by the synergistic effects of p-Tau and glucose hypometabolism. This observation suggests that additional non-necroptotic cell death pathways, or mechanisms independent of the p-Tau-RIPK1 interaction, may compensate for or contribute to neuronal loss under these conditions and warrant further investigation.

## Supporting information

SupplementalL Figure titles and legends, Method

## RESOURCE AVAILABILITY

### Lead contact

Further information and requests for resources and reagents should be directed to and will be fulfilled by the lead contact, Chengyu Zou.

### Materials availability

All mouse lines are available from the lead contact with a completed Materials Transfer Agreement.

### Data and code availability

All data reported in this paper will be shared by the lead contact upon request. This paper does not report original code. Any additional information required to reanalyze the data reported in this paper is available from the lead contact upon request.

## ACKNOWLEDGMENTS

We thank Dr. Zhuohao He for his valuable advice during the preparation of this manuscript. We thank the staff members of the Large-scale Protein Preparation System (https://cstr.cn/31129.02.NFPS.LSPS) at the National Facility for Protein Science in Shanghai (https://cstr.cn/31129.02.NFPS), for providing technical support and assistance in data collection and analysis. We also thank the human brain donors and their families for facilitating this research and the staff members of the Neurobiobank Munich for their preserving work. This study was supported in part by the National Natural Science Foundation (NSF) of China (82188101, 22425704, and 82170234), Shanghai Municipal Science and Technology Major Project (22JC1410400), Shanghai Municipal Science and Technology Major Project (2025), Shanghai Basic Research Pioneer Project, Shanghai Key Laboratory of Aging Studies (19DZ2260400), CAS Pioneer Initiative (2024000026), National Special plan for High-level Talents (2022000178), Young Talent Cultivation Program of CAS at Shanghai (2023000010) Guangzhou Planned Project of Science and Technology (2024A03J1178),Chinese Academy of Science, Shanghai Branch (Grant No. JCYJ-SHFY-2022-005), the CAS Project for Young Scientists in Basic Research (Grant No.YSBR-095), the Strategic Priority Research Program of the Chinese Academy of Sciences (Grant No. XDB1060000). Dr. Cong Liu is a SANS Exploration Scholar. Dr. Matthias Brendel was funded by the Deutsche Forschungsgemeinschaft (DFG) under Germany’s Excellence Strategy within the framework of the Munich Cluster for Systems Neurology (EXC 2145 SyNergy, ID 390857198).

## AUTHOR CONTRIBUTIONS

Conceptualization, X.C., S.L., J.Y., C.L., J.H. and C.Z.; methodology, X.C., S.L., A.N., X.L., M.B., M.Z., B.S., and C.Y.; writing—original draft, X.C. and C.Z.; writing—review & editing, X.C., J.Y., C.L., J.H. and C.Z.; funding acquisition, C.L., J.H. and C.Z.; resources, B.S., J.Y., C.L., J.H. and C.Z.; supervision, C.L., J.H. and C.Z.

## SUPPLEMENTAL INFORMATION

Figures S1–S9, Table S1-S3 and Video S1–S2

