## SupplementalL Figure titles and legends, Method for "Glucose hypometabolism and hyperphosphorylated Tau synergistically drive neuronal necroptosis"

### SUPPLEMENTAL FIGURE TITLES AND LEGENDS

#### Figure S1. related to Figure1.

(A) Immunoblotting of Tau and p-Tau protein levels in Ctrl HT22, WT-Tau HT22 and P301S-Tau HT22 cells. AT8 antibody was used for the detection of p-Tau.

(B) Immunoblotting of Tau aggregates marker in WT-Tau HT22 and P301S-Tau HT22 cells exposed to low glucose exposure (LGE, 0.7mM glucose in culture medium) for the indicated times.

(C) Immunoblotting of apoptosis and pyroptosis markers in Ctrl HT22 and P301S-Tau HT22 cells exposed to LGE for the indicated time periods.

(D-E) WT-Tau HT22 and P301S-Tau HT22 cells were cultured in medium containing 0.7 mM glucose for the indicated times. Cell death was assessed by SYTOX Green (SG) positivity assay (D). The levels of indicated proteins were determined by immunoblotting (E).

(F) Immunoblotting of AMPK pathway markers in Ctrl HT22, WT-Tau HT22 and P301S-Tau HT22 cells exposed to LGE for the indicated time periods.

(G) P301S-Tau HT22 cells were cultured in DMEM containing 0.7 mM glucose with or without the TNF-α blocking antibody (2 μg/mL) for the indicated durations. Cell death was assessed by SG staining.

(H) P301S-Tau HT22 cells were cultured in high glucose DMEM and then treated with TNF-α (100 ng/mL), 5Z-7-oxozeaenol (5Z-7, 0.25 μM), z-VAD-fmk (20 μM), with or without the TNF-α blocking antibody (2 μg/mL), for the indicated times. Cell death was assessed by SG staining.

(I) Representative images of primary cortical neurons isolated from WT and PS19 mice exposed to culture medium containing 1.5 mM or 0.7 mM glucose for 36 hours, with the treatment of Ferr-1 (10 μM), z-VAD (20 μM), CQ (25 μM) and DTT (2 mM). Immunofluorescence staining for MAP2 (green) and AT8 (magenta) is shown. Scale bar is 50 μm.

(J) Quantifications of MAP2-positive neuron counts and AT8 staining intensity as shown in (I). Quantified data are presented as mean ± SEM. Unpaired two-tailed Student's t test was used. *$p < 0.05$, **$p < 0.01$, n.s = not significant.

#### Figure S2. related to Figure2.

(A) Immunoblotting of A20 expression in primary cortical neurons isolated from WT and PS19 mice exposed to low glucose exposure (LGE, 0.7mM glucose in culture medium) for the indicated times.

(B) Flow cytometry gating strategy of NeuN-positive cells in the brain.

(C) Quantification of A20 expression in NeuN-positive cells using flow cytometry. n = 5 mice/group.

(D) Immunoblotting of A20 expression in hippocampal lysates from MLKL-shRNA AAV injected mice.

(E-F) Representative images of Sudan Black staining in serial brain sections from PS19 mice injected with AAVs encoding either an empty vector (Ctrl) or MLKL shRNA (E). Quantification of lateral ventricle and hippocampus area is shown (F). n = 7 mice/group. Scale bar, 1 mm.

(G-H) Immunofluorescence analysis of pyramidal cell layer thickness via NeuN staining in the CA1 region of PS19 mice injected with AAVs encoding either empty vector (Ctrl) or MLKL shRNA. n = 7 mice/group. Scale bar is 50 μm.

(I-K) Cognitive behavior analysis of AAV injected PS19 mice. (I) Novel arm preference, quantified as the percentage of time spent in the novel arm, during the T-maze test. (J) Escape latency during training trials and (K) target quadrant preference, quantified as the percentage of time spent in the target quadrant, during the probe test of Morris water maze. n = 8 mice/group.

(L) Quantification of swimming speed in PS19 mice injected with AAVs encoding either an empty vector (Ctrl) or MLKL shRNA during the Morris water maze test. Quantified data are presented as mean ± SEM. Unpaired Student's t test was used (F, H and K) and One sample t test against 50% was used (I). *p < 0.05, **p < 0.01, n.s = not significant.

### Figure S3. related to Figure2.

(A) Immunoblotting of A20 expression in hippocampal lysates from A20-shRNA AAV injected mice.

(B-E) Representative images and quantification of pRIPK1 (B-C) and pMLKL (D-E) immunostaining in the CA1 region of WT, PS19 and PS19xD138N mice, three weeks after injection with A20-shRNA or Ctrl AAV. Colocalization of pRIPK1 or pMLKL (green, indicated by red arrowheads) with NSE (gray) and Dapi (blue) are shown. n = 7 mice/group. Scale bar is 10 μm.

(F) Immunoblotting of A20 expression in primary cortical neurons isolated from WT and PS19 mice exposed to LGE for the indicated times, with or without ALCAR (20 mM).

(G) A20 levels in P301S-Tau HT22 cells under the indicated treatments were confirmed by immunoblotting.

(H) P301S-Tau HT22 cells transfected with lentivirus carrying A20 shRNA or control shRNA were cultured in medium containing 0.7 mM glucose. Cell death was assessed using an SYTOX Green (SG) positivity assay. Where indicated, ALCAR (20 mM), Nec-1s (10 μM) or GSK-872 (1 μM) was added to the culture medium.

(I) Ctrl HT22 cells transfected with lentivirus carrying A20 shRNA or control shRNA were cultured in medium containing 0.7 mM glucose. Cell death was assessed using an SG positivity assay. Quantified data are presented as mean ± SEM. Unpaired two-tailed Student's t test was used. *p < 0.05, n.s = not significant.

**Figure S4. related to Figure3.**

(A-B) Representative images (A) and quantification (B) of A20 immunostaining in microglia within the CA1 region of ALCAR treated PS19 mice. A20 (gray), IBA1 (green), Dapi (blue) staining are shown. n = 6 mice/group. Scale bar, 20 μm.

(C) Quantification of swimming speed in ALCAR-treated PS19 mice during the Morris water maze test. Quantified data are presented as mean ± SEM. One-way ANOVA followed by Tukey's multiple comparisons test was used. $^{*}p < 0.05$, $^{**}p < 0.01$, n.s = not significant.

**Figure S5. related to Figure4.**

(A) Quantification of A20 positivity and pRIPK1 positivity in hippocampal CA regions and the subiculum of three AD patients who underwent FDG-PET scanning during lifetime.

(B) Representative images and quantification of HE immunostaining in hippocampal subregions of AD patients. Scale bars are 1 mm (overview) and 50 μm (magnified view).

(C) Quantification of the density of NFT-positive neurons in hippocampal CA regions and the subiculum of AD patients. n = 10 patients. Quantified data are presented as mean ± SEM. A paired two-tailed t-test was used (A) and unpaired Student's t test was used (B and C). $^{*}p < 0.05$.

**Figure S6. related to Figure5.**

(A) HEK 293 cells were transfected for 48 hours with plasmids encoding HA-tagged P301S Tau and Flag-tagged RIPK1 truncation mutants (as indicated). Cell lysates were subjected to immunoprecipitation using anti-HA-conjugated beads. Interactions between P301S Tau and RIPK1 truncations were analyzed by immunoblotting.

(B) HEK 293 cells were co-transfected with expression plasmids encoding HA-tagged P301S Tau and Flag-tagged RIPK1-ID truncations (as indicated) containing a C terminal GFP tag. Cell lysates were subjected to immunoprecipitation using anti-HA-conjugated beads, and the immunoprecipitates were analyzed by immunoblotting.

(C) Immunoblot analysis of phosphorylated Tau levels in recombinant Tau protein purified from E. coli and subjected to in vitro phosphorylation by p38, with phosphorylation detected using AT8 antibody.

(D) Immunoblot analysis of phosphorylated Tau levels in recombinant Tau protein purified from E. coli and subjected to in vitro phosphorylation by MARK2-T208E, with phosphorylation detected using p-Tau (Ser 416) antibody.

**Figure S7. related to Figure5.**

(A) RMSD curves show the root-mean-square deviation (RMSD) of the peptide over the course of the simulation (right). Peptide sequences and phosphorylation sites used for molecular dynamics simulations are shown in the left panel.

(B) Energy decomposition of key residues in RIPK1-CRS.

(C) Residues at the binding interface between p-Tau and RIPK1-CRS in the stable conformation. Interface residues on p-Tau are shown in red, and those on RIPK1-CRS are shown in blue.

(D) HEK 293 cells were co-transfected with expression plasmids encoding HA-tagged P301S Tau and Flag-tagged RIPK1 mutants (as indicated). Cell lysates were subjected to immunoprecipitation using anti-HA-conjugated beads, and the immunoprecipitates were analyzed by immunoblotting.

**Figure S8. related to Figure7.**

(A) Immunoblotting of RIPK1 protein levels in RIPK1 KO MEFs transfected with lentivirus to stably express either full-length RIPK1 or RIPK1-△CRS along with P301S Tau.

(B) RIPK1 KO MEFs were cotransfected with lentivirus to stably express either full-length RIPK1 or RIPK1-△CRS, along with P301S Tau. Cells were treated with TNF-α (10 ng/ml), 5Z-7 (0.25 μM), with or without z-VAD-fmk (20 μM), for the indicated times. TNF-induced apoptosis and necroptosis were assessed using MTT assay and SYTOX Green (SG) staining, respectively.

(C) Representative images of primary cortical neurons isolated from WT and PS19 mice. Neurons were first transduced with lentivirus expressing RIPK1 CRS or empty vector (Ctrl) for 48 hours, and then exposed to culture medium containing 1.5 mM or 0.7 mM glucose for 36 hours. Immunofluorescence staining for MAP2 (green) and AT8 (magenta) is shown.

(D) Quantifications of MAP2-positive neuron counts and AT8 staining intensity as shown in (C). Quantified data are presented as mean ± SEM. Unpaired two-tailed Student's t test was used. $^{**}p < 0.01$, n.s = not significant.

**Figure S9. related to Figure7.**

(A-C) Immunostaining images and quantification of hippocampal sections from PS19 mice injected with AAVs encoding the mCherry-tagged RIPK1 CRS domain. Neuronal (NeuN, green), microglial (IBA1, green), and astrocytic (GFAP, green) markers were visualized alongside mCherry (red) and nuclear counterstaining with DAPI (blue). Scale bars are 200 μm (overview) and 20 μm (magnified view). Quantification of colocalization for the indicated signals is shown in the right panels.

(D-F) Immunofluorescence imaging of hippocampal sections from PS19 mice without AAV injection. Neuronal (NeuN, green), microglial (IBA1, green), and astrocytic (GFAP, green) markers were visualized alongside mCherry (red) and nuclear counterstaining with DAPI (blue). Scale bars are 200 μm (overview) and 20 μm (magnified view).

(G) Quantification of swimming speed during the Morris water maze test in PS19 mice injected with AAVs encoding either an empty vector (Ctrl) or the RIPK1 CRS domain. Quantified data are presented as mean ± SEM. One-way ANOVA followed by Tukey's multiple comparisons test was used. $^{**}p < 0.01$, n.s = not significant.

**Table S1**

Demographic and clinical characteristics of postmortem human brain samples.

**Table S2**

List of proteins identified by mass spectrometry as interacting partners of p-Tau.

**Table S3**

Quantitative analysis of Western blot data throughout the manuscript.

**Video S1**

The dynamic interaction between the RIPK1 CRS domain and Tau-F was visualized by MD stimulations.

**Video S2**

The dynamic interaction between the RIPK1 CRS domain and p-Tau-F was visualized by MD stimulations.

### EXPERIMENTAL MODEL AND STUDY PARTICIPANT DETAILS

#### Human samples

Human tissue was obtained from the Neurobiobank Munich. Brain donations were made following informed consent and collected in accordance with the guidelines of the local ethics committee at Ludwig-Maximilians University Munich, as well as the Code of Conduct of BrainNet Europe[67]. The use of tissue was approved by the Neurobiobank Munich committee. Ten Alzheimer's disease cases, balanced for sex and classified as Braak and Braak stage V or VI, were selected for analysis. The mean age at death was 75.6 years (range: 56.9-86 years) (Table S1).

Hippocampal sections of formalin fixed and paraffin embedded tissue were stained with Hematoxylin and Eosin, and immunohistochemical diaminobenzidine stainings were performed. We applied the monoclonal antibody AT8 (ThermoFisher, #MN1020) for phosphorylated Tau, the monoclonal antibody clone 4G8 (BioLegend, #800711) for Amyloid beta, the monoclonal Anti-TNFAIP3 antibody [EPR2663] (abcam, ab92324) for A20, and the monoclonal Anti-pRIPK1 (Ser166) (D8I3A) antibody (Cell Signaling, #44590) for phosphorylated RIPK1.

After digitizing the stainings with a Zeiss Axio Scan Z.1 scanner with a magnification of 20 (pixel size: 0.22 * 0.22 µm²), hippocampal subregions were manually annotated in Qupath (version 0.5.1)[68]. Three, if space-wise not possible two, representative tiles of 500 * 500 µm² were labeled for each subregion, i.e., CA1, CA2, CA3, CA4 and subiculum, and the mean of the value of interest was calculated for each region, respectively. Subsequently, CA1 to CA4 were combined by calculating the mean across these subregions. Neurons were counted manually based on the nucleus and soma morphology in Hematoxylin and Eosin stainings. Amyloid beta, p-Tau (AT8), A20, A20 positive neurites, and pRIPK1 positivity were evaluated semiquantitatively in four levels (0: negative, 1: sparse, 2: moderate, 3: frequent positive staining) for each tile separately. Additionally, neurons with pRIPK1 positive soma and neurofibrillary tangle bearing neurons were counted to determine the ratio between them.

Three out of ten AD patients received fluorodeoxyglucose Positron Emission Tomography scans (FDG-PET) and MRI during their lifetime. Hippocampal subregions were visually identified in axial and coronal views of structural MRIs of these patients. Based on this region segmentation, the FDG PET signal was collected for CA1, CA2-4, and subiculum of the brain hemisphere that passed postmortem immunohistochemical analysis. Signal intensities were controlled for individual cerebellar gray matter values. In parallel to the histological quantification, the mean of the CA regions was calculated (CA1-4 = (3*CA2-4 + CA1)/4) resulting in one FDG-PET value for CA1-4 and one for subiculum, respectively. FDG-PET images were acquired on a Siemens ECAT EXACT HR+ PET scanner. All patients had fasted for at least six hours, and had a maximum plasma glucose level of 150 mg/dl at time of scanning. A single intravenous dose of 140 ± 7 MBq FDG was administered while the patients rested in a room with dimmed light and low noise level, where they remained undisturbed for 20 minutes. After positioning in the scanner, emission was recorded from 30 to 60 min p.i.. PET data were reconstructed with filtered-back-projection.

#### Mice

Heterozygous transgenic mice (Tg(Prnp-MAPT*P301S)Ps19Vle/JNju) were kindly provided by Prof. Zhuohao He at IRCBC. Age-matched non-transgenic littermates were used as wild-type (WT) control mice. PS19 mice were crossed with *Ripk1*$^{D138N/D138N}$ mice to generate PS19xD138N mice[69]. All mice were maintained under pathogen-free conditions and housed with no more than five animals per cage under standardized light-dark cycle conditions with ad libitum access to food and water. The vivarium was maintained under controlled temperature (21 ± 1°C) and humidity (50 - 60%). Only male mice were used in this study to eliminate sex-related variability, particularly in behavioral testing, where female mice may exhibit increased variability due to hormonal cycles and estrous-related behavioral fluctuations. To minimize variability, all treatment comparisons were performed strictly between littermates throughout the study. The experimental procedures used in this study were approved by the Ethics Committees of Interdisciplinary Research Center on Biology and Chemistry (IRCBC), Chinese Academy of Sciences.

**Cell lines and primary cell cultures**

HEK293T, HT22 and MEF cells were cultured and passaged in high glucose Dulbecco's modified Eagle medium (DMEM, Gibco) with 10% (vol/vol) fetal bovine serum (FBS, Gibco) and 100 U/ml penicillin/streptomycin. The P301S-Tau HT22, WT-Tau HT22, and control HT22 cell lines were generated using pMSCV-HA-P301S Tau, pMSCV-HA-WT Tau, or empty pMSCV vectors, respectively, with lentiviral supernatants produced in HEK293T cells, followed by puromycin selection to establish stable lines. Primary cortical and hippocampal neurons were dissected from pregnant PS19 mice at embryonic day 17 according to the established protocol[70]. Primary neurons were maintained in Neurobasal Medium (Gibco) supplemented with 1% 100X GlutaMAX™ (Gibco), 2% B27 (Gibco) and 1% penicillin/streptomycin. All cells were cultured at 37 °C with 5% $CO_2$. To adjust glucose concentrations as indicated, glucose-free DMEM or Neurobasal Medium was used in combination with supplemental glucose solution.

**Cell death quantification**

Cell death quantification was performed as previously described[71]. Cells were seeded into 384-well plates one day prior to analysis. On the day of the experiment, cells were washed three times with PBS to remove residual culture medium and then incubated in culture medium containing the indicated glucose concentrations, with or without specified compounds, in the presence of 1 mM Sytox Green (Invitrogen). Sytox Green fluorescence was measured at specified time points. Cell death was quantified using the formula: (Detected fluorescence − background fluorescence) / (Maximum fluorescence − background fluorescence) × 100. Maximum fluorescence was determined by fully permeabilizing the cells with 0.1% Triton X-100.

**Immunoblotting**

The cells were harvested and lysed using NP40 buffer. The NP40 lysis buffer contained 50 mM Tris (pH 7.5), 130 mM NaCl, 1 mM Na3VO4, 50 mM NaF, 50 mM NEM, 1 mM PMSF, and a 1x protease inhibitor cocktail. The protein concentration in the cell supernatants was measured using a BCA assay and adjusted to ensure equal loading. The proteins were then separated by SDS-PAGE and subjected to immunoblotting with the specified antibodies.

**Immunoprecipitation**

Immunoprecipitation were performed referring to the procedures described previously[72]. Cell lysates were prepared in NP40 lysis buffer containing protease inhibitors and then centrifuged at 12,000 rpm for 10 minutes. The protein concentration in the supernatants was determined using a BCA assay (Thermo), and the concentrations were equalized across samples. For immunoprecipitation, 1 mL of each supernatant was transferred to a new 1.5 mL microcentrifuge tube, and 20 µL of Anti-HA Affinity Beads (Smart-lifesciences) or Anti-Flag Affinity Beads (Smart-lifesciences) were added to each tube and incubated at 4 °C for at least 4 hours. After incubation, the beads were collected and washed three times with cold lysis buffer. The washed beads were then boiled with 80 µL of loading buffer for 5 minutes at 95 °C. The boiled samples were centrifuged at 1,000 g for 5 minutes, and the supernatants containing proteins were loaded onto an SDS-PAGE gel. The proteins were subsequently analyzed by immunoblotting with indicated antibodies.

**Immunohistochemistry**

Mice were perfused with cold PBS. The brain was then dissected and immersed in 4% paraformaldehyde for 24 hours at 4°C. After fixation, brains were affixed to the base and then sectioned into 50 µm slices. The brain sections were blocked for 2 hours at room temperature with a solution of PBS containing 0.1% Triton X-100 and 5% normal goat serum (NGS). After blocking, the sections were incubated with primary antibodies overnight at 4°C. The following day, the sections were thoroughly washed and then incubated with secondary fluorescent antibodies at room temperature for 2 hours. The expression of A20 was analyzed using Manders' colocalization coefficients calculated with the JACoP plugin in ImageJ.

**Immunofluorescence**

For immunostaining, cells were grown on coverslips in a 12-well plate. After being washed with PBS, the cells were fixed in 4% paraformaldehyde for 15 minutes at room temperature. Following fixation, the cells were permeabilized with 0.2% Triton X-100 in PBS for 15 minutes and blocked with 5% goat serum in PBST (0.2% Triton X-100 in PBS) for 1 hour. Next, the cells were incubated with primary antibodies overnight at 4 °C, followed by the incubation with secondary antibodies at room temperature for 1 hour. After being washed three times with PBS, the cells were mounted on glass slides for imaging.

**TurboID proximity labelling**

TurboID proximity labelling protocol was adopted from a previous study[73]. HT22 cells were transfected with expression vectors for TurboID-tagged P301S Tau for 48 hours. After LGE treatment, biotin was added to culture medium to a final concentration of 500 µM and incubated for 20 minutes at 37 °C. Reactions were halted by aspirating the medium, placing the cells on ice, and performing three consecutive 1-minute washes with ice-cold 1× PBS. Cells were then processed as detailed for immunoprecipitation, with the following changes. Protein lysates in NP40+Urea buffer (25 mM Tris-HCl pH 7.5, 135 mM NaCl, 0.5% NP-40, 1 mM EDTA, 1 mM EGTA, 10 mM β-glycerol phosphate, 5% glycerol, 6 M Urea and 2% SDS supplemented with protease inhibitors) were incubated with 20 µL of streptavidin agarose beads (GE Life Sciences) overnight at 4 °C on a rotating wheel. The beads were then washed thrice in NP40 buffer, five times in dd$H_2O$. Affinity-purified biotinylated proteins were processed for on-bead digestion and mass spectrometry analysis as described below.

**Mass spectrometry and data analysis**

Affinity-purified biotinylated proteins were trypsin digested on-beads. The resulting peptides in three replicates were analyzed on a a nanoElute LC system coupled to a timsTOF Pro mass spectrometer (Bruker, Bremen, Germany). Peptide identification and quantification were performed by FragPipe v22[74]. The tandem mass spectra were searched against the UniProt mouse protein database supplemented with a set of frequent contaminants. The precursor mass tolerance was set as 20 ppm and the fragment mass tolerance was set as 0.1 Da. The cysteine carbamidomethylation was set as a static modification and the methionine oxidation was set as a variable modification. The false discovery rates at the peptide spectral match and protein levels were both controlled at <1%. The unique and razor peptides were used for protein quantification. Protein quantification was performed using unique and razor peptides, and MaxLFQ intensities were used for statistical analysis[75]. Missing values were imputed using the Perseus-style imputation method implemented in FragPipe-Analyst.

.

**Lentivirus construction and cell transfection**

To produce lentiviruses encoding A20 shRNA (ATTCGATGAAACATAGAGTGC), control shRNA (GTCTCCACGCGCAGTACATTT), A20, P301S-Tau, WT-Tau, the RIPK1 CRS domain, or empty vector controls, HEK 293T cells were transfected accordingly. Viral supernatants were harvested 48 hours post-transfection and filtered through a 0.22 µm membrane. HT22 cells were then infected with the filtered supernatants in the presence of 10 µg/mL polybrene. Following infection, the cells were maintained in culture for an additional two days before puromycin selection and downstream applications.

**Protein expression and purification**

The Tau, GFP-tagged RIPK1 CRS and GFP plasmids were transformed into BL21 (DE3) Chemically Competent Cell (EC1002M, Shanghai Weidi Biotechnology). Cells were grown to an $OD_{600}$ of 0.6 and induced with 1 mM IPTG overnight at 16 °C. Proteins were loaded onto Ni Smart beads with buffer containing 50 mM Tris-HCl, pH 8.0, 500 mM NaCl, and 10% glycerol. The proteins were eluted with imidazole and then further purified using a size exclusion column. The Tau, GFP-RIPK1 CRS and GFP proteins were purified using a Superdex 200 16/600 column (GE Healthcare). The purified proteins were stored in a buffer containing 50 mM Tris-HCl, pH 8.0, 500 mM NaCl, and 10% glycerol at −80 °C.

**In vitro phosphorylation of Tau**

Tau was phosphorylated in vitro by either p38 or MARK2-T208E, following previously described protocols[44,45]. p38 mediated reaction were carried out at 30°C in 15 µL volumes containing 8.8 µg of Tau, 3 mM ATP, 10 mM $MgCl_2$, 50 mM Tris-HCl (pH 7.5), 1 mM ethylene glycol tetraacetic acid (EGTA), 1 mM dithiothreitol (DTT), 1 µM okadaic acid, 1 mM sodium vanadate, 1 mM benzamidine-HCl, 5 µg/mL leupeptin, 2 µg/mL aprotinin, 1 µg/mL pepstatin, 0.5 mM PMSF, and 0.36 µg of active p38. The reactions were terminated after 7.5 hours by adding 5 µL of 4× Laemmli sample buffer (Laemmli, 1970), followed by heating at 100 °C for 5 minutes and storage at −20°C until further analysis. For MARK2-mediated Tau phosphorylation, Tau was incubated with cat MARK2-T208E at a molar ratio of 10:1 in a buffer of 50 mM HEPES, pH 8.0, 150 mM KCl, 10 mM $MgCl_2$, 5 mM EGTA, 1 mM PMSF, 1 mM DTT, 2 mM ATP (Sigma), and protease inhibitor cocktail (Roche) at 30 degrees overnight.

**Surface Plasmon Resonance**

Biacore 8K analyses were carried out on an NTA sensor chip. The system was primed with HBS-EP$^+$ running buffer (10 mM HEPES, 150 mM NaCl, 3 mM EDTA, 0.05% P20, pH 7.4). The NTA surface was charged by injecting 0.5 mM $NiCl_2$ at 5-10 µL/minute for 1-2 minutes. His-Tau or His-p-Tau (10 µg/mL in running buffer) was then injected at 5-10 µL/minute for 1-3 minutes, yielding an immobilization level of ~50-200 resonance units; the surface was washed to stabilize the baseline. Serial dilutions of RIPK1 CRS (in running buffer) were injected at 30 µL/minute for 1-5 minutes, followed by a 2-10 minutes dissociation phase. The NTA surface was regenerated by injecting 350 mM EDTA or 10-100 mM imidazole for 30-60 seconds and then recharged with $NiCl_2$ for subsequent cycles. Sensorgrams were analyzed with Biacore 8K Evaluation Software to derive the equilibrium dissociation constant ($K_D$).

**Molecular dynamics simulations**

The structures of Tau, phosphorylated Tau (p-Tau), and the RIPK1-CRS domain were predicted using AlphaFold3. To improve structural stability and resolve potential steric clashes, Rosetta Relax was employed for energy minimization. The relaxed structures were saved in PDB format for downstream docking and simulation analyses. Protein-protein docking was performed using the local version of HDOCK. This platform utilizes a hierarchical fast Fourier transform (FFT)-based algorithm to rapidly sample potential binding conformations in both translational and rotational space. Each docked pose was scored based on a composite function that integrates electrostatic interactions, hydrogen bonding, van der Waals forces, and hydrophobic effects. The top-ranked conformation was selected for subsequent molecular dynamics (MD) simulations.

All-atom MD simulations were conducted using Gromacs 2023.3[76]. The AMBER force field was applied to describe peptide systems[77,78]. Hydrogen atoms were added using the pdb2gmx utility. Each protein or complex was solvated in a truncated cubic box filled with TIP3P water molecules, maintaining a minimum distance of 10 Å between solute and box edge. To mimic physiological conditions, $Na^+$ and $Cl^-$ ions were added to a final concentration of 0.145 M. Topology and parameter files were generated accordingly.

Energy minimization of the system was first performed using the steepest descent algorithm (mdrun), with an initial step size of 0.01 nm and a maximum force threshold of 1000 kJ/mol • nm. Equilibration was performed in two phases. First, a 100-ps NVT (constant volume and temperature) simulation was carried out to gradually heat the system from 0 K to 310.15 K, ensuring even solvent distribution. Second, a 100-ps NPT (constant pressure and temperature) simulation was conducted using the Berendsen barostat to equilibrate system pressure to 1 bar. Subsequently, production MD simulations were run for 50 ns. All bonds involving hydrogen atoms were constrained using the LINCS algorithm. A 2-fs time step was used throughout the simulations. Electrostatic interactions were computed using the particle-mesh Ewald (PME) method with a real-space cutoff of 1.2 nm. Nonbonded interactions were truncated at 10 Å, and the neighbor list was updated every 10 steps. Post-simulation, trajectory files were corrected for periodic boundary conditions and analyzed for root-mean-square deviation (RMSD) and other structural parameters.

**Fluorescence Imaging of Liquid-Liquid Phase Separation**

The purified proteins were diluted in a buffer containing 50 mM Tris (pH 7.5) to a final concentration of 50 μM for Tau and p-Tau (fluorescent labeled with Alexa Fluor™ 568 C5 maleimide), or 5 μM for GFP-tagged RIPK1 CRS and GFP. Phase separation (PS) of Tau, p-Tau, GFP-tagged RIPK1 CRS, and GFP was induced by adding 5% (w/v) PEG. After phase separation occurred, 3 μL of the solution was pipetted onto a glass slide for confocal imaging. The images were collected using a Leica TCS SP8 microscope with a 100x objective (oil immersion, NA = 1.4) at room temperature.

**Fluorescence recovery after photobleaching (FRAP) assay**

The FRAP assay was executed using the FRAP module on a Leica TCS SP8 confocal microscope, equipped with a 100× oil immersion objective. This procedure involved selectively bleaching fluorescently labeled assemblies with a laser beam and targeting a specific circular region of interest. Following photobleaching, imaging was performed continuously, capturing one frame every 20 seconds. The fluorescence intensity in the bleached region (Itm) was measured, along with the intensity (Itc) in a nearby unbleached assembly serving as a control. For quantitative analysis, the fluorescence intensity at the bleached site at each time point (t) was normalized against the control. The recovery of fluorescence was calculated using the formula: It = (Itm/I0m)/(Itc/I0c). All captured images were subsequently analyzed using the Leica Application Suite X software.

**Brain tissue protein extraction**

Mouse brain tissue was homogenized in lysis buffer (25 mM Tris-HCl pH 7.5, 135 mM NaCl, 0.5% NP-40, 1 mM EDTA, 1 mM EGTA, 10 mM β-glycerol phosphate, 5% glycerol and 1% TritonX-100 and 0.1% SDS with phosphatase and protease inhibitors). Homogenates were incubated on ice for 1 hour and centrifuged at 12,000g for 10 minutes at 4 °C. The protein concentration in the cell supernatants was measured using a BCA assay and adjusted to ensure equal loading. The proteins were then separated by SDS-PAGE and subjected to immunoblotting with the specified antibodies.

**Magnetic resonance imaging**

Magnetic resonance imaging (MRI) was performed referring to the procedures described previously[79]. MRI was performed on a 9.4T small-animal MRI system (Bruker BioSpec, Germany). Experimental mice were anesthetized with a 1.5% isoflurane/oxygen mixture throughout the imaging process. T2-weighted images were scanned. The MRI scanning sequences and parameters are as follows: TR (repeat time) = 3200 ms, TE (echo time) = 33 ms, field of view (scan field) = 13.26 x 8.84 mm, matrix (matrix dimensions) = 156 x 104, SI (layer spacing) = 0.5 mm, FA (reverse Angle) = 180°, slices (layers) = 20. ITK-SNAP software was used to reconstruct the hippocampus in 3D and calculate the hippocampus and ventricle volumes.

**Stereotaxic Injection and viral vectors**

For in vivo viral injection experiments, mice were anesthetized with 0.5% isoflurane mixed with 1% $O_2$. All AAV viral vectors used in this study were packaged as adeno-associated virus serotype 9 (AAV9) and driven by the human synapsin (hSyn) promoter to ensure neuron-specific expression in vivo. Virus ($1\times10^{12}$ viral genomes/ml) were stereotaxically injected into the hippocampus of both hemispheres at a dose of 1μL. The following coordination was used: -2.92 mm to bregma, ± 3.00 mm from midline, -3.45 mm from dura and -1.58 mm to bregma, ± 1.50 mm from midline, -1.80 mm from dura. The injection was done with a 10-μL microsyringe at a speed of 0.5 μL/min for the first 0.1 μL and 0.1 μL/min for the rest. Control mice were bilaterally injected with empty vector virus (1 μL) with the same equipment and speed. After recovery from the anesthetization, mice were transferred back to their home cages with regular housing conditions until the next experiment.

For A20 knockdown, the experimental virus PFD-rAAV-hSyn-EGFP-5′miR-30a-shRNA(A20)-3′-miR30a-WPREs (shA20 sequence: ATTCGATGAAACATAGAGTGC) was used together with the control virus rAAV-hSyn-EGFP-5′miR-30a-shRNA(scramble)-3′-miR-30a-WPRE. For the RIPK CRS overexpression group, the experimental vector rAAV-hSyn-RIPK1 CRS-P2A-mCherry-WPRE was employed, with rAAV-hSyn-mCherry-WPRE-hGHpolyA serving as the control virus. For the MLKL knockdown experiments, the experimental virus rAAV-hSyn-EGFP-5′miR-30a-shRNA(mlkl)-3′-miR-30a (shMLKL sequence: CATTGGAATACCGTTTCAG-AT) and its corresponding control rAAV-hSyn-EGFP-5′miR-30a-shRNA(scramble)-3′-miR-30a were used.

**Novel T maze**

Novel T maze was conducted according to a previously described[36,80]. The goal of the Novel T Maze is to determine if a mouse can use spatial cues to distinguish a novel arm from a familiar arm that it had previously visited. Each mouse run twice in the T-Maze, first the 3 minutes Habituation phase then a 2 minutes Test phase. The delay or intertrial interval (ITI) between the end of the habituation phase and start of the test phase is 2 minutes. During the habituation phase, one arm (either left or right) is blocked off. The start arm always remains the same. Test analyses show time of the Novel Arm versus Familiar Arm.

**Morris water maze**

The Morris water maze was conducted as previously described[81]. Morris water maze test was used to evaluate the spatial learning and memory ability of mice. The water maze is conducted in a circular tank with a radius of 60 cm. Titanium dioxide was added to water before use. A hidden platform was placed in a selected target quadrant during the training phase. Mice were placed into the maze randomly from one of the four directions and were allowed to look for the platform for 60 seconds. The mice that could not find the platform were guided to the platform for 15 seconds. Four training sessions were conducted per day for each mouse. On day 5, the hidden platform was removed, and mice were subjected to a probe test. Escape latency to find the platform and time spent in the target quadrant during the probe test were recorded and analyzed.

### QUANTIFICATION AND STATISTICAL ANALYSIS

Data were analyzed using GraphPad Prism. A two-tailed Student's t test was used for pairwise comparison between two groups. For multiple comparisons within three or more conditions, we performed one-way ANOVA followed by Tukey's multiple comparisons. *$p < 0.05$, **$p < 0.01$, n.s = no significant difference. Differences were significant if p value < 0.05 (*). Results are presented as mean ± SEM unless further specified. All behavioral tests, image acquisition, image analysis, and cell death analysis were performed blinded with respect to mouse genotype, treatment condition and human sample information.
