## Supplementary figures and images for "Glucose hypometabolism and hyperphosphorylated Tau synergistically drive neuronal necroptosis"

### Supplemental Figure

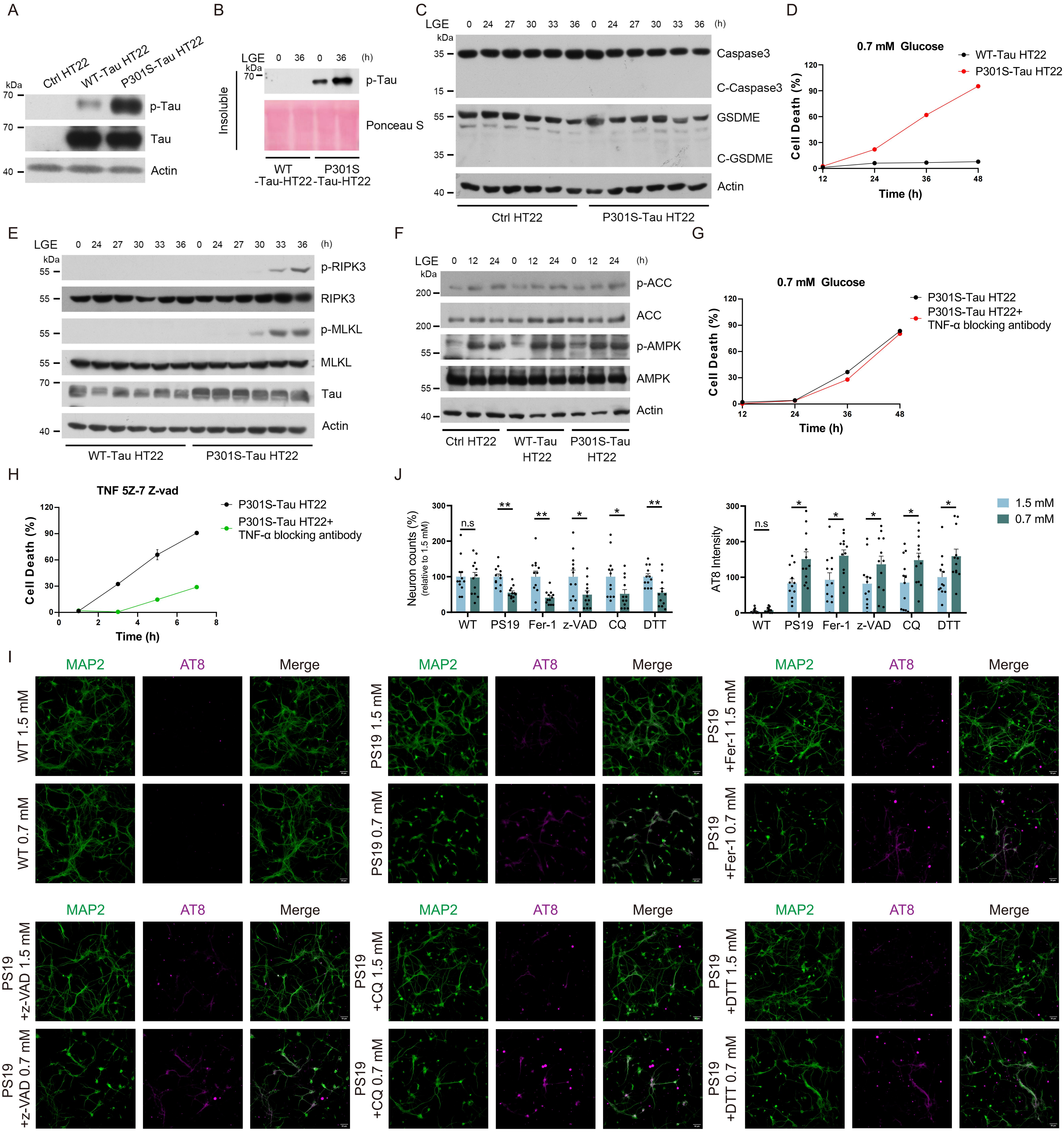

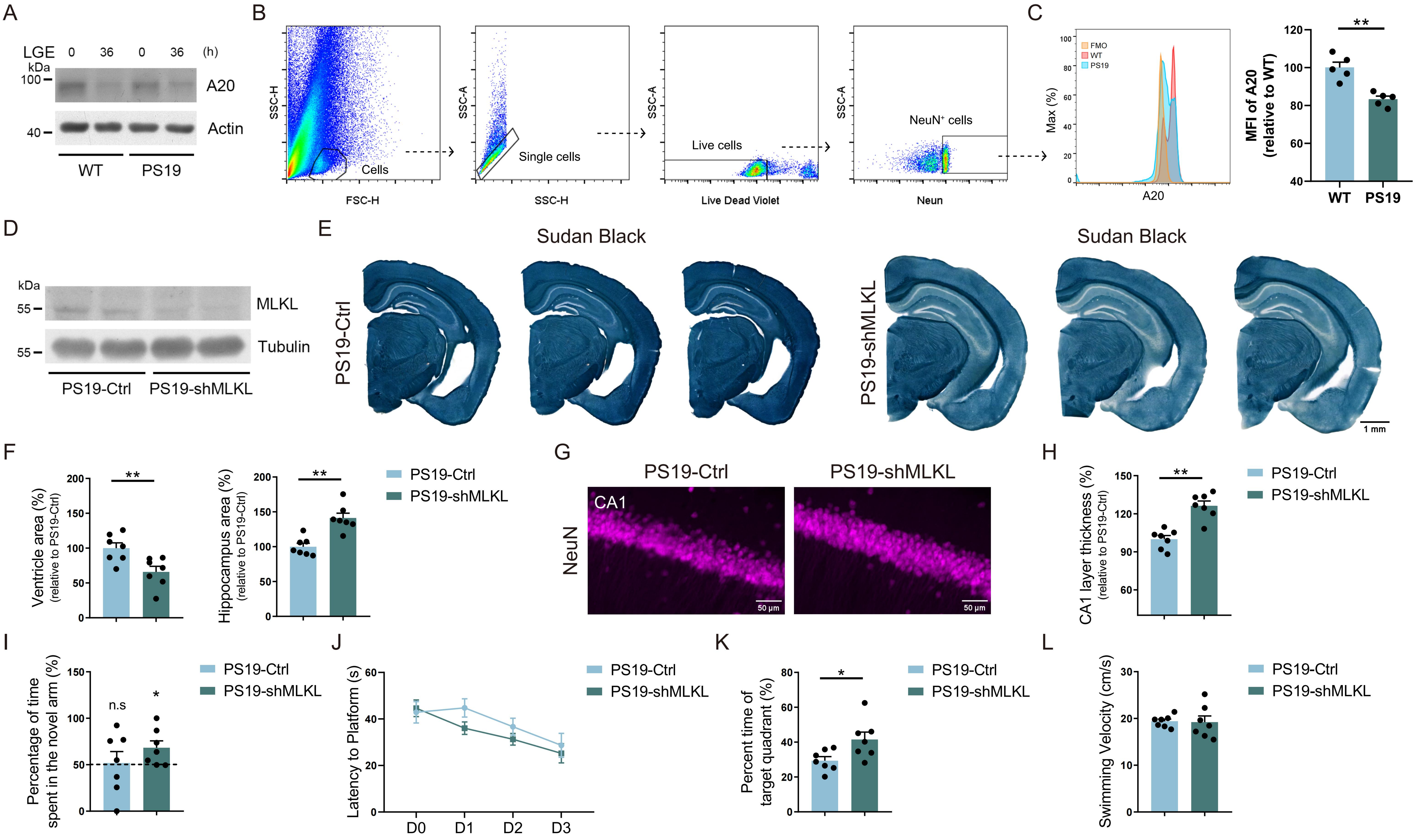

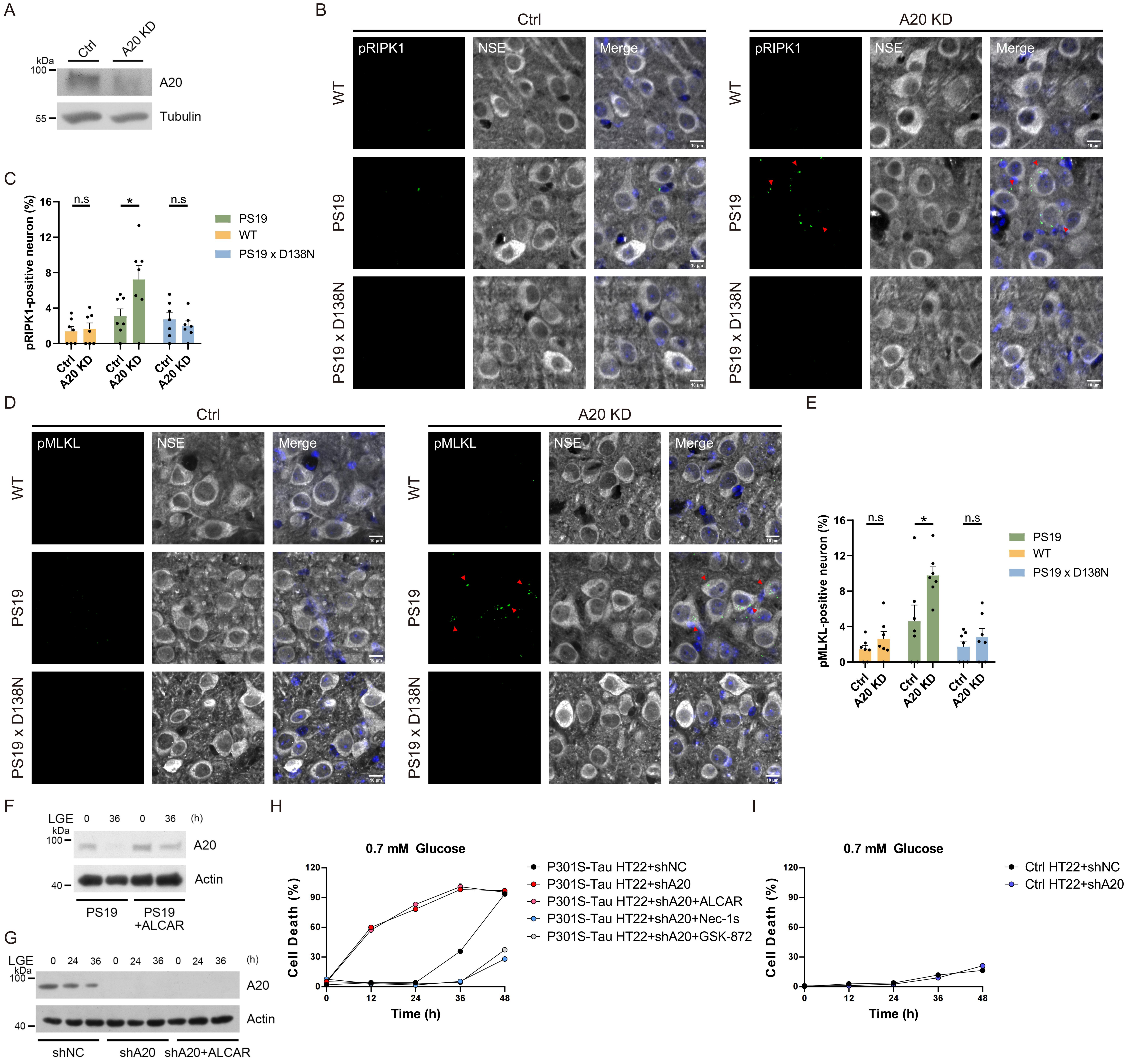

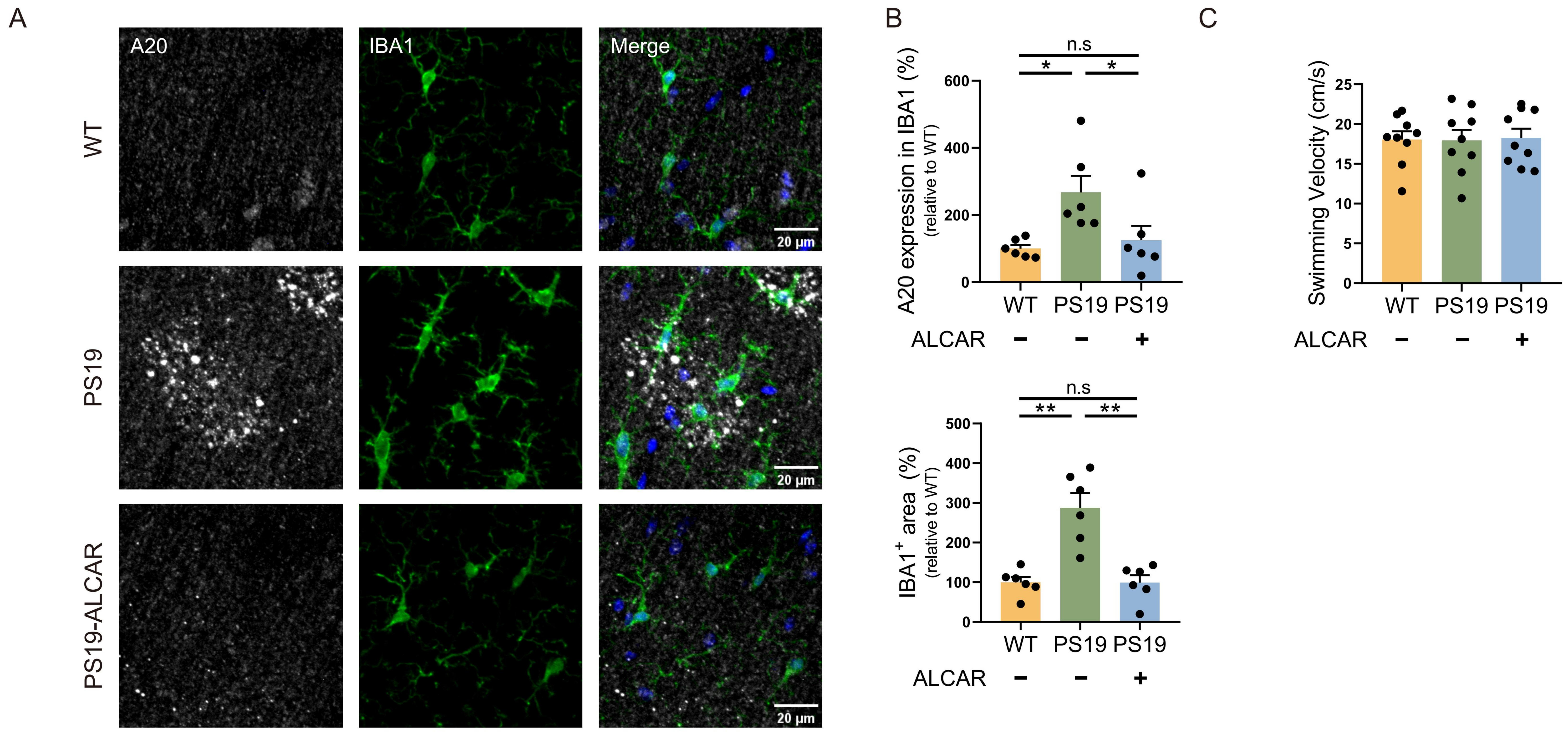

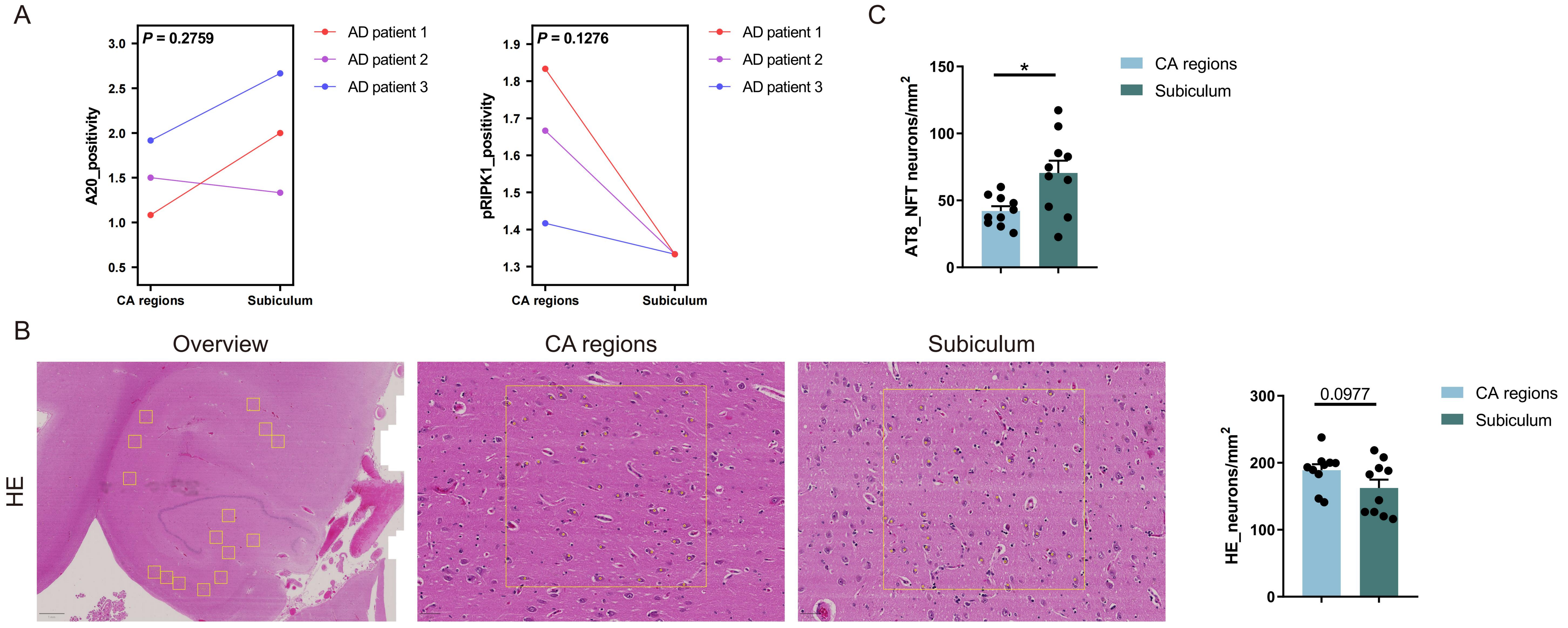

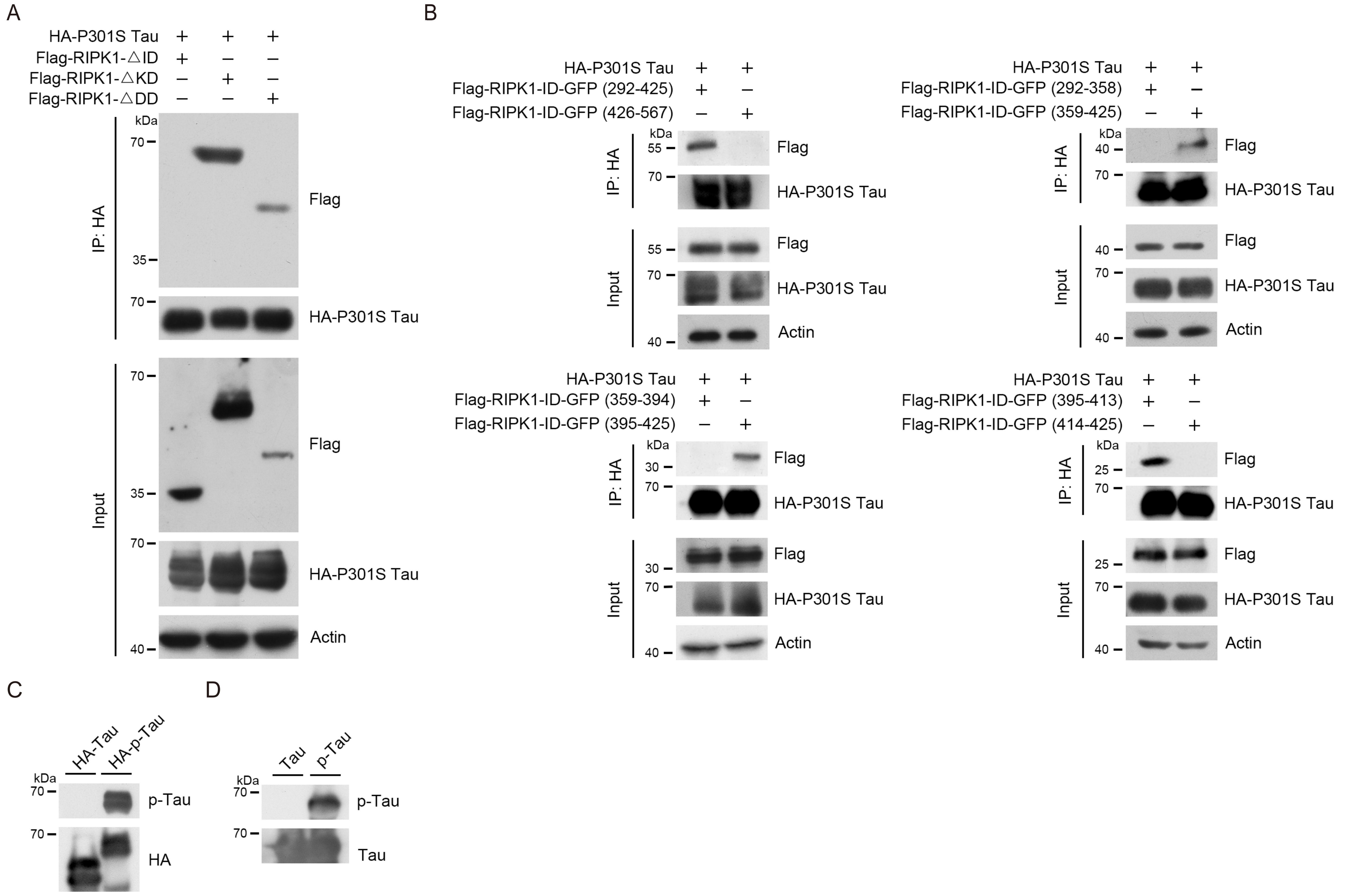

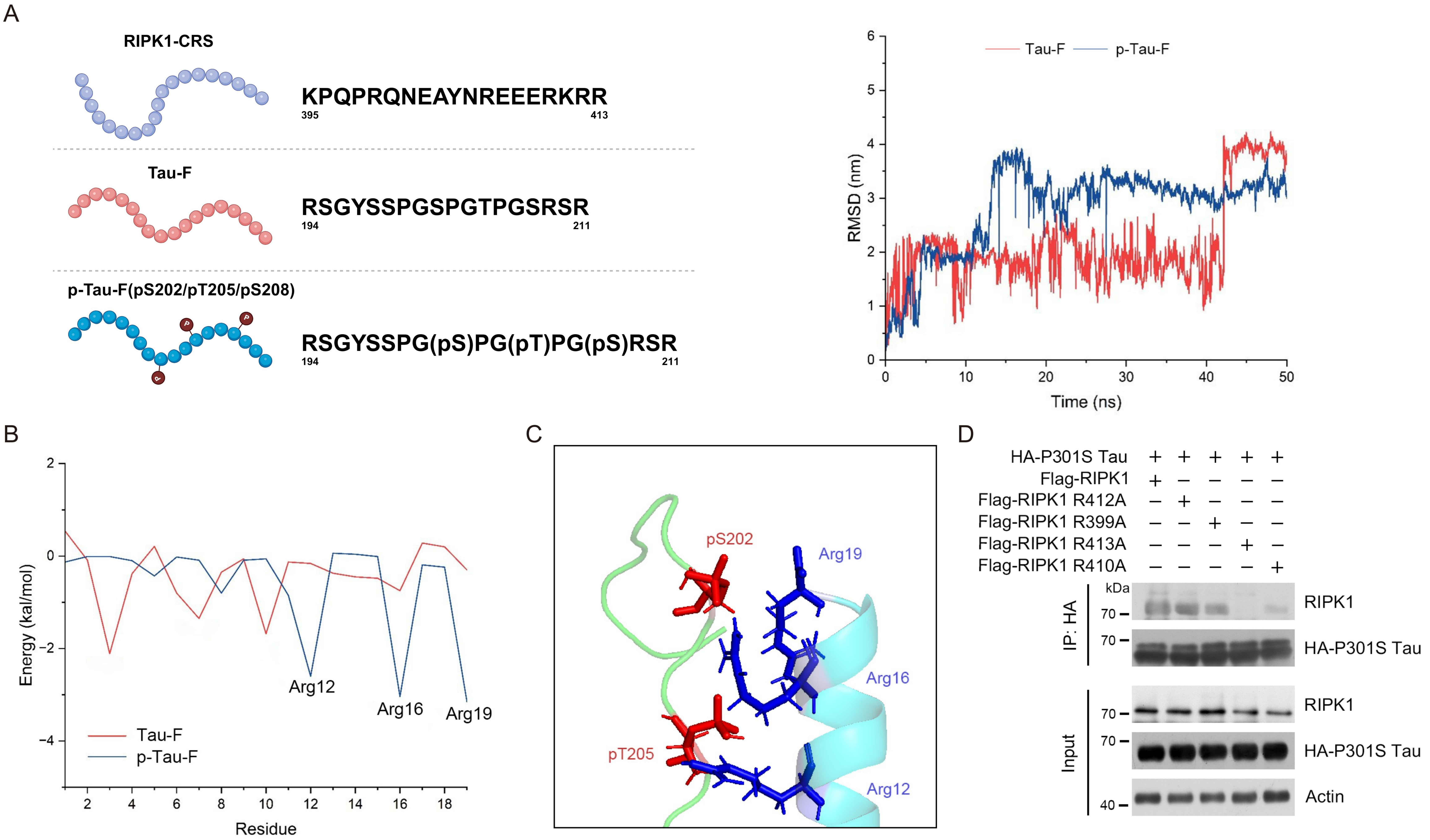

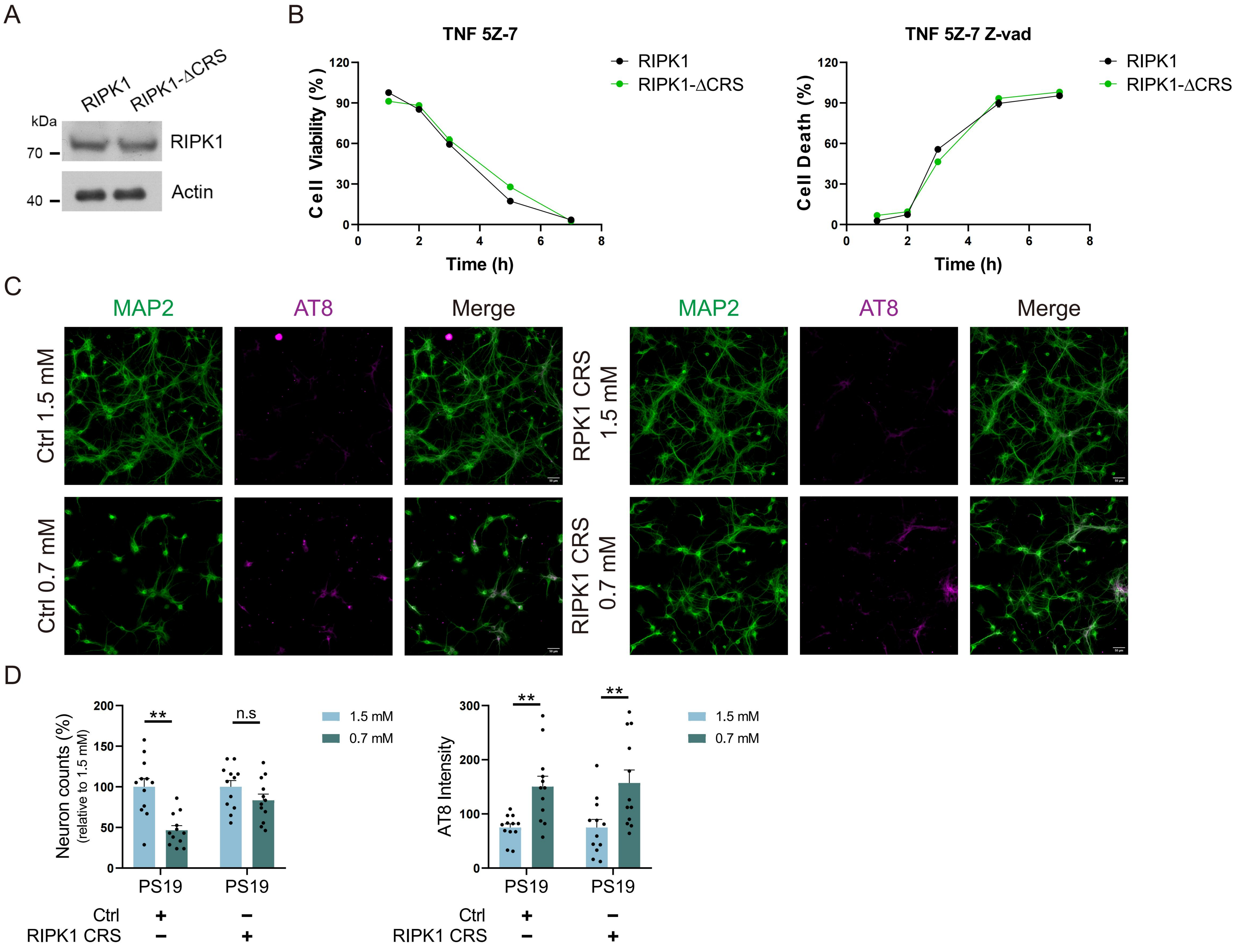

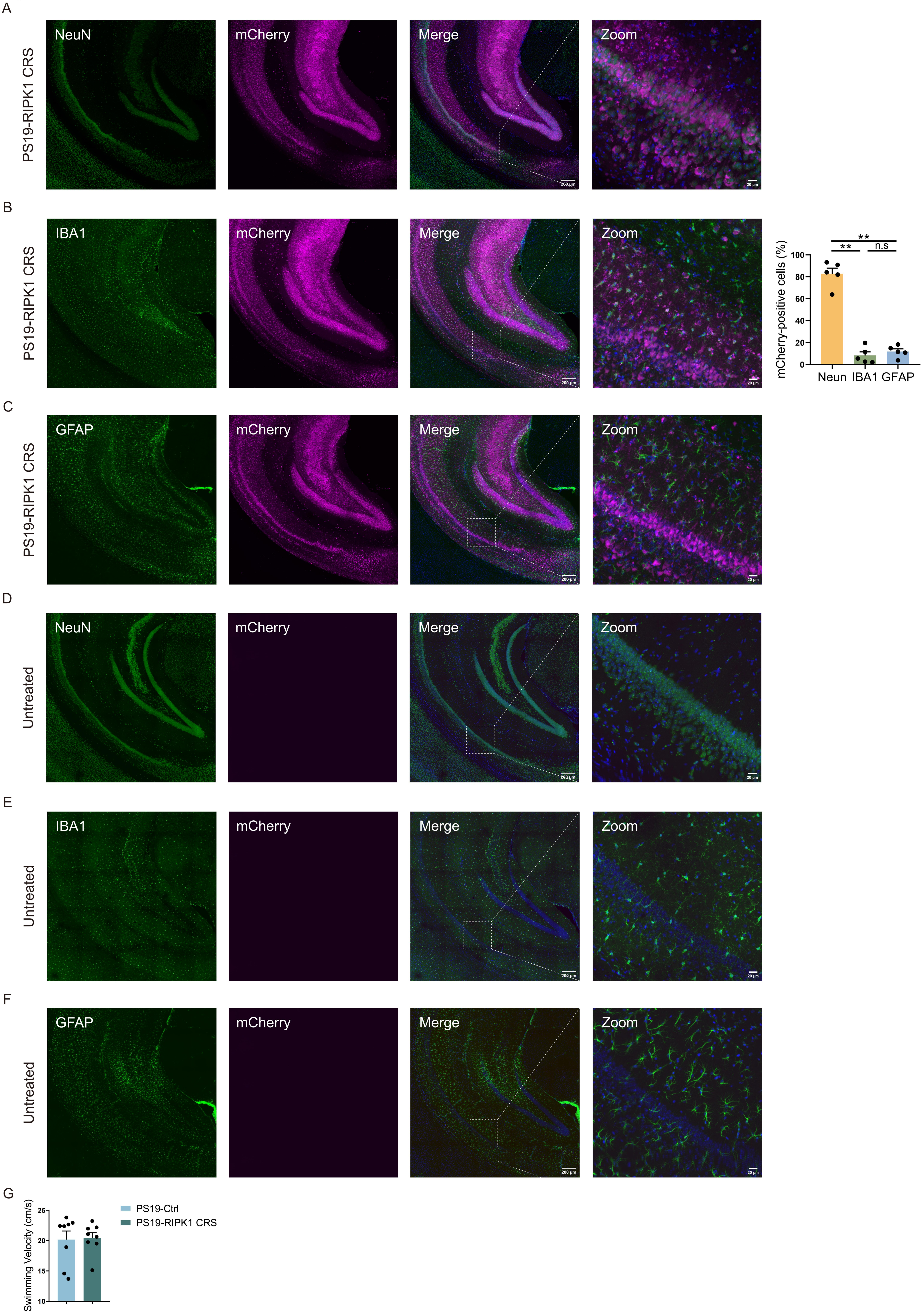
